# 14-3-3 binding maintains the Parkinson’s associated kinase LRRK2 in an inactive state

**DOI:** 10.1101/2024.11.22.624879

**Authors:** Juliana A. Martinez Fiesco, Ning Li, Astrid Alvarez de la Cruz, Riley D. Metcalfe, Alexandra Beilina, Mark R. Cookson, Ping Zhang

**Affiliations:** Kinase Complexes Section, Center for Structural Biology, Center for Cancer Research, National Cancer Institute, Frederick, MD, 21702, USA; Cell Biology and Gene Expression Section, National Institute on Aging, National Institutes of Health, Bethesda, MD, 20892, USA

## Abstract

Leucine-rich repeat kinase 2 (LRRK2) is a central player in cellular signaling and a significant contributor to Parkinson’s disease (PD) pathogenesis. 14-3-3 proteins are essential regulators of LRRK2, modulating its activity. Here, we present the cryo-electron microscopy structure of the LRRK2:14-3-3_2_ autoinhibitory complex, showing that a 14-3-3 dimer stabilizes an autoinhibited LRRK2 monomer by binding to key phosphorylation sites and the COR-A and COR-B subdomains within the Roc-COR GTPase domain of LRRK2. This interaction locks LRRK2 in an inactive conformation, restricting LRR domain mobility and preventing dimerization and oligomer formation. Our mutagenesis studies reveal that PD-associated mutations at the COR:14-3-3 interface and within the GTPase domain reduce 14-3-3 binding, diminishing its inhibitory effect on LRRK2. These findings provide a structural basis for understanding how LRRK2 likely remains dormant within cells, illuminate aspects of critical PD biomarkers, and suggest therapeutic strategies to enhance LRRK2-14-3-3 interactions to treat PD and related disorders.

## Introduction

Mutations enhancing leucine-rich repeat kinase 2 (LRRK2) activity are a leading cause of familial PD and genetic variation at the same locus significantly contributes to lifetime risk of sporadic PD^1–7^. LRRK2 is a large multidomain protein of 2527 residues, possessing both GTPase and kinase domains^8–14^. Its catalytic core comprises a Roco family GTPase domain, including a GTP-binding Ras of complex proteins (Roc) domain coupled with a C-terminal of Roc domain (COR, split into COR-A and COR-B subdomains), and a serine/threonine kinase domain. This core is flanked by N-terminal armadillo (ARM), ankyrin (ANK), and leucine-rich repeats (LRR) domains, and a C- terminal WD40 domain (Fig. 1a)^15–19^. LRRK2 plays crucial roles in the endolysosomal system, notably through the phosphorylation of specific Rab proteins^20–23^. Pathogenic mutations associated with PD, located in the Roc, COR, and kinase domains (Supplementary Fig. 1a), are known to increase kinase activity and/or decrease GTPase activity^1–7,24–33^. Elevated LRRK2 kinase activity has also been associated with an increased risk of cancer^34–38^.

**Fig. 1.**
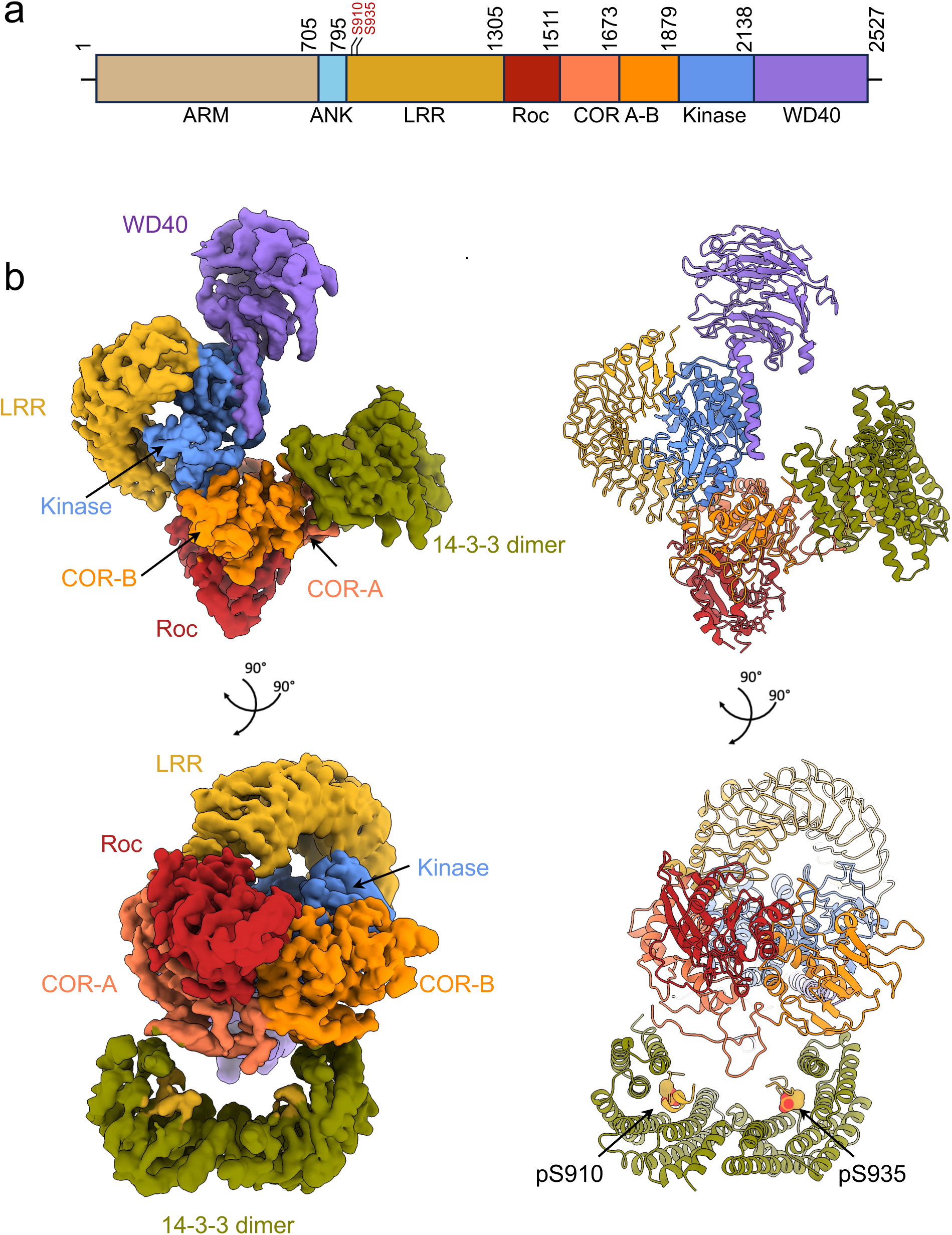
Structure of the LRRK2:14-3-3_2_ complex. **a.** Schematic representation of LRRK2 domain organization. Residues S910 and S935 are in red, which serve as 14-3- 3 binding sites upon phosphorylation. **b.** Cryo-EM density map at a resolution of 3.96 Å (left), with the corresponding structural model (right) colored as in **(a)** and shown in two different orientations for clarity.

The molecular mechanisms driving LRRK2 kinase activation are poorly understood; however, factors such as phosphorylation^39,40^, oligomerization^41–50^, membrane association^51–56^, and complex formation with regulatory proteins are all potentially important contributors^57–60^. Recent data from several laboratories indicates that LRRK2 is relatively inactive in cells unless triggered by damage to lysosomes, which then leads to the accumulation of phosphorylated RAB proteins on membranes^51–56^. Members of the 14-3-3 family are established LRRK2 interactors and are proposed to contribute to LRRK2 stability ^61,62^. 14-3-3 are regulatory proteins ubiquitously expressed and abundantly present in cells, acting as dimeric scaffolds^63,64^ modulate a broad spectrum of client proteins through various mechanisms^65–70^. Several PD-associated mutations such as R1441C/G/H, Y1699C, and I2020T, have diminished 14-3-3 interaction, which is correlated with increased kinase activity^71–73^. Additionally, studies on PD rodent models and analyses of postmortem PD brains show reduced LRRK2 and 14-3-3 interactions, associated with increased kinase activity in sporadic PD^74^. Collectively, these results indicate that 14-3-3 binding is inhibitory for LRRK2 activity.

14-3-3 proteins are phosphoserine/phosphothreonine-binding proteins^75^. Recent structural studies have elucidated the interactions of 14-3-3 proteins with essential kinases and scaffold proteins^76–79^. In LRRK2, the N-terminus features a loop preceding the LRR domain that includes a cluster of potential 14-3-3 binding sites^61,80^. Additional potential 14-3-3 binding sites have also been suggested within the Roc domain and the C-terminus of the protein (Supplementary Fig. 1a)^81^. Despite the robust interaction between 14-3-3 proteins and LRRK2, the exact molecular details, such as binding sites, stoichiometry, and the impact on LRRK2’s oligomerization and kinase activity, remain underexplored. This hinders a comprehensive mechanistic understanding of the role of 14-3-3 proteins in maintaining cytosolic LRRK2 in a physiological inactive state and prevents an understanding of how disrupted LRRK2/14-3-3 interactions contribute to PD pathogenesis.

Here, we report the cryo-EM structure of the full-length monomeric LRRK2 complexed with a 14-3-3 dimer, revealing how critical phosphorylation sites and GTPase subdomains engage in the inactive state of LRRK2. Our findings provide the structural basis for how 14-3-3 is able to modulate LRRK2 kinase activity and oligomerization state. Our insights into the LRRK2 and 14-3-3 interactions will enable future development of therapeutic strategies aimed at modulating this interaction.

## Results

### Formation and structural characterization of the LRRK2:14-3-3_2_ complex

To study the interaction between LRRK2 and 14-3-3 proteins, we expressed and purified both proteins separately, utilizing the monomeric form of LRRK2 and 14-3-3 gamma (γ), the most abundant 14-3-3 isoform in the brain^82^ and with the highest affinity for LRRK2-derived peptides^81^ (see Methods, Supplementary Fig. 1). We successfully formed the LRRK2/14-3-3 complex under conditions optimized for ionic strength (Supplementary Fig. 2), enabling subsequent structural analysis. Mass spectrometry and mass photometry analyses confirmed the presence of both proteins in the complex (Supplementary Fig. 2). Initial cryo-electron microscopy (cryo-EM) studies revealed 2D classes and a 3D density map consistent with the LRRK2/14-3-3 complex formation (Supplementary Fig. 3). However, the density corresponding to 14-3-3 was poorly defined, with a substantial population of LRRK2 not bound to 14-3-3. To enhance the complex homogeneity, we applied cross-linking with bis(sulfosuccinimidyl)suberate (BS3) prior to size-exclusion chromatography (Supplementary Fig. 4), which improved the density of 14-3-3 within the complex.

Further three-dimensional reconstruction of the cross-linked particles yielded a 3.96 Å map with a density corresponding to a LRRK2 monomer and additional density matching a 14-3-3 dimer, confirming the LRRK2/14-3-3 complex formation (Fig. 1b, Supplementary Figs. 5 and 6). The structural details, including visible α-helices and β- strands, were well resolved, consistent with a map reconstructed at this resolution (Supplementary Fig. 6). The complex demonstrated a 1:1 stoichiometry between a LRRK2 monomer and a 14-3-3 dimer (Fig. 1b), referred to as the LRRK2:14-3-3_2_ complex.

Within the complex, LRRK2 adopted a conformation similar to the previously reported inactive LRRK2 monomer^45^ (Cα root mean square deviation (RMSD) of 0.4 Å, Supplementary Fig. 7a). The N-terminal region (residues 1-906), including the ARM and ANK domains, was absent from the density map, suggesting that they are flexible.

The elongated LRR domain covered the kinase domain, occluding and preventing the recruitment of protein substrates to the kinase domain. Although the kinase domain was nucleotide free, density within the Roc domain suggested the presence of bound GDP. Additionally, the conformation of the switch I loop in the Roc domain corroborated this GDP-bound state (Supplementary Fig. 6e). The 14-3-3 dimer, characterized by nine antiparallel α-helices per protomer and forming a cup-like shape with two client-binding grooves^70^ is positioned adjacent to the catalytic core of LRRK2, making contacts with the COR domain. Local refinement for the LRRK2:14-3-3_2_ interacting region, specifically the Roc-COR:14-3-3_2_ part, modestly enhanced the resolution for this region to 3.87 Å (Supplementary Fig. 6b).

Interestingly, despite using purified recombinant 14-3-3γ for the LRRK2:14-3-3_2_ complex formation, mass spectrometry detected multiple 14-3-3 isoforms in the sample, (Supplementary Fig. 2b). Given the highly conserved client-interacting residues across all 14-3-3 isoforms (Supplementary Fig. 7b), it is likely that endogenous 14-3-3 proteins co-purified with the LRRK2 sample. This suggested that LRRK2:14-3-3 complex forms and is stable *in vivo*. To investigate this hypothesis further, we co-expressed both proteins in mammalian cells and successfully purified the complex, showing the *in vivo* formation of the LRRK2 /14-3-3 complex (Supplementary Fig. 8).

### 14-3-3 dimer binds to pS910 and pS935 sites and the COR domain in LRRK2

The interactions of 14-3-3 proteins with client proteins are usually mediated through primary and secondary interactions between the client and 14-3-3. Primary interactions are established when phospho-serine/threonine-containing motifs of the client proteins bind to the conserved amphipathic groove of 14-3-3, formed by the α-3, α-5, α-7, and α- 9 helices of 14-3-3^83^. Secondary interactions, less commonly observed, involve larger interfaces between the globular domain of the client protein and additional surface on 14-3-3^83^, enhancing the specificity and stability of the complex. Identifying the primary binding sites on LRRK2 has been challenging due to its lack of conventional binding motifs^84,85^. A primary sequence analysis of LRRK2 does not provide clear 14-3-3 binding sites (Supplementary Fig. 9a). Despite this, experimental data have demonstrated that LRRK2 possesses several highly phosphorylated serine residues located between the ANK and LRR domains, specifically S910, S935, S955, and S973^41,61,86–88^, raising questions about complex formation and stoichiometry between LRRK2 and 14-3-3.

The LRRK2:14-3-32 structure clearly revealed that a 14-3-3 dimer binds LRRK2 at pS910 and pS935 sites as primary interactions (Fig. 2). The cryo-EM map showed substantial densities for the 14-3-3 binding motifs across residues 907-919 and 930- 940, bound to each protomer of the 14-3-3 dimer, with clear density for the phosphate groups at pS910 and pS935 (Fig. 2a). This finding was consistent with our mass spectrometry data, which indicated that both S910 and S935 sites were highly phosphorylated (Supplementary Fig. 9b). Previously noted for its high flexibility in other LRRK2 structures^45,50,89^, this region exhibited significant stabilization upon 14-3-3 binding. The phosphate groups at S910 and S935 were positioned to interact with the positively charged groove of 14-3-3, including the canonical R57, R132, and Y133 triad as well as K50 within the α-3 helix (Fig. 2b). Additionally, extensive hydrophobic interactions between LRRK2 residues 911-919 and 936-940 and the 14-3-3 binding groove were established. The loop region (residues 920-929) connecting the binding sites in each 14-3-3 protomer is dynamic with lower resolution. Although this region becomes visible when contouring the density map at a low threshold, it did not provide sufficient detail to enable model building (Supplementary Fig. 10a). Interestingly, this loop contains residue Q923, and a LRRK2 Q923H mutation has been reported in one Brazilian patient with positive family history of PD^90^. Similarly, the region between residue 942 and 983 which connects these primary interaction motifs with the LRR was flexible and not visible on the map (Supplementary Fig. 10a). These primary interactions in the complex were similar to those observed in the crystal structure of human 14-3-3 in complexes with phospho-peptides containing the prominent phosphorylation sites pS910 and pS935^91^.

**Fig. 2.**
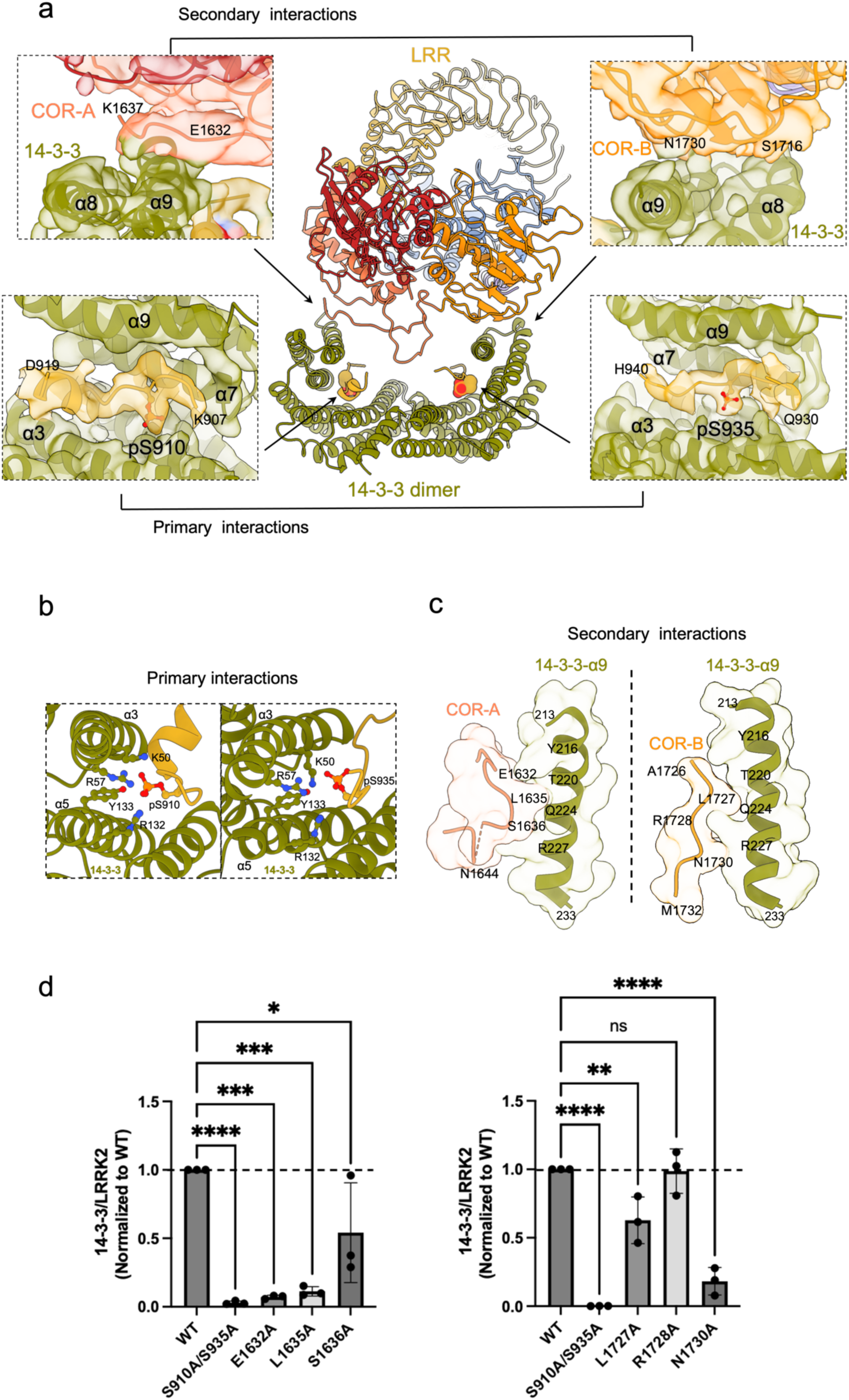
Detailed interactions and mutational effects at the LRRK2:14-3-3_2_ binding interfaces. **a.** Overview of the LRRK2:14-3-3_2_ contact regions in LRRK2:14-3-3_2_ complex. Insets detail the primary and secondary interaction sites, supported by the corresponding cryo-EM densities. **b.** Close-up view of the primary interactions, with LRRK2 phosphorylation sites, pS910 and pS935, engaging with the 14-3-3 substrate binding grooves. **c.** Close-up view of the secondary interactions with LRRK2 COR-A and COR-B subdomain residues contacting the α-9 helices of the 14-3-3 dimer. **d.** Quantitative analysis of LRRK2/14-3-3 interactions through Co-IP experiments of LRRK2 with endogenous 14-3-3 comparing WT LRRK2 with mutants in the COR-A (left) and COR-B (right) at the LRRK2:14-3-3 secondary interface, as well as primary interface mutants S910/S935A. Data illustrate the impact of mutations on the interaction strength. Refer to Supplementary Fig. 11 for representative membrane images and source data for complete membrane images. Data are mean ± SEM (n = 3 independent experiments), significance of difference was quantified using one-way Brown-Forsythe and Welch ANOVA test and reported in the source data file.

Mutation of S910 and S935 to alanine abolished the binding between LRRK2 and 14-3- 3, as demonstrated in our co-immunoprecipitation (Co-IP) experiments using Flag-tagged LRRK2 and endogenous 14-3-3 (Fig. 2d). This emphasized the critical role of these residues in forming the primary interactions with 14-3-3 and explained the dependency of the LRRK2:14-3-3 interaction on ionic strength of the solution in vitro (Supplementary Fig. 2a). Affinity studies with short peptides containing LRRK2 sequences suggested that residues at the Roc domain (S1444) and C-terminus (T2524) of LRRK2 are part of potential primary binding motifs for 14-3-3^81^. However, in our LRRK2:14-3-32 structure, these sites are occluded by LRRK2 intramolecular interactions (Supplementary Fig. 10b) and were not solvent exposed to establish interactions with 14-3-3.

Additionally, our LRRK2:14-3-3_2_ structure revealed a unique, previously unidentified interface between the COR domain and the 14-3-3 dimer, which served as the secondary interactions between LRRK2 and 14-3-3. Residues within 1632-1644 and 1727-1732 regions in the COR-A and COR-B subdomains were aligned along the interfaces and interacted with both α-9 helices in both monomers in the 14-3-3 dimer (Fig. 2), covering buried surface areas of 186 and 278 Å^2^, respectively. This secondary interaction was established mainly through Van der Waals interactions. Specifically, LRRK2 COR-A residues E1632, L1635, and S1636 and COR-B residues L1727, R1728, and N1730 at the secondary interfaces make numerous contacts with 14-3-3 α-9 helices residues Y216, Q224, and R227 in the model (Fig. 2c). We took a mutational approach to determine whether these COR residues also contributed to the overall interaction between the two proteins. In the Co-IP assay using LRRK2 with those residues mutated to alanine, we observed a reduction in binding of ∼40 - 90%, with mutations E1632A, L1635A, and N1730A displaying the largest effect (Fig. 2d), demonstrating that while the segment containing the pS910/S935 establish the primary and strongest interaction with 14-3-3 dimer, the secondary interactions between 14-3-3 and COR-A and COR-B subdomains also contribute to the overall stability of the interaction between the two proteins.

In addition, and consistent with previous studies^61,62,71,72^, LRRK2 activity was inhibited by 14-3-3. We further demonstrated that the modulation of these interfaces affected the inhibition of LRRK2 kinase by 14-3-3. The secondary interface observed here, located outside the 14-3-3 cradle, is unusual, however, similar interfaces have been observed in the structures of other 14-3-3/client interactions (for example, BRAF/14-3-3 and Exoenzyme T/14-3-3 complexes^79,92^). The residues involved in the primary interaction with LRRK2 are highly conserved among the different 14-3-3 isoforms (Supplementary Fig. 7b), and similarly, those involved in the secondary interface are conserved in all human 14-3-3 family members, indicating that a similar interface would be expected regardless of the isoform composition of the 14-3-3 dimer (Supplementary Fig. 7b).

COR domains are well-documented dimerization modules in Roco proteins^17^and have been shown to facilitate the assembly of LRRK2 into various oligomeric forms under specific experimental conditions, including dimers, tetramers, and higher-order oligomers^45,50,93^. These oligomeric structures, if physiologically relevant, are significant for understanding both the function and pathological implications of LRRK2^94^. Our structural analysis revealed a partial overlap between the COR:14-3-3 interface and the LRRK2 interface in the inactive homodimer, mediated by the COR-B subdomains from each LRRK2 protomer (Fig. 3a). Specifically, residues 1727-1730 in COR-B, crucial for the LRRK2 homodimer interface, directly interact with 14-3-3 in the LRRK2:14-3-3_2_ complex (Fig. 3b). We investigated if 14-3-3 could bind LRRK2 homodimers by conducting a series of multi-angle light scattering (MALS) experiments with pre-formed LRRK2 dimer and increasing concentrations of recombinant 14-3-3γ (Fig. 3c, Supplementary Fig. 11). We observed no disruption of the LRRK2 dimer with increasing 14-3-3γ concentration, at higher concentrations, the measured mass (643 KDa) was consistent with the binding of a single 14-3-3γ dimer to a single LRRK2 dimer. However, the overlap in the interface may result in reduced dimerization affinity, so 14-3-3 binding may competitively inhibit LRRK2 dimerization. 14-3-3 binding may also prevent the formation of higher molecular states, such as the LRRK2 tetramer observed in cryo-EM grids, which consists of two inactive and two active LRRK2 monomers, due to interface overlap (Supplementary Fig. 10d). Additionally, this interaction would also be expected to impede the association between LRRK2 and microtubules, where it forms microtubule-bound filaments through COR-COR interactions in a closed active conformation^93^.

**Fig. 3.**
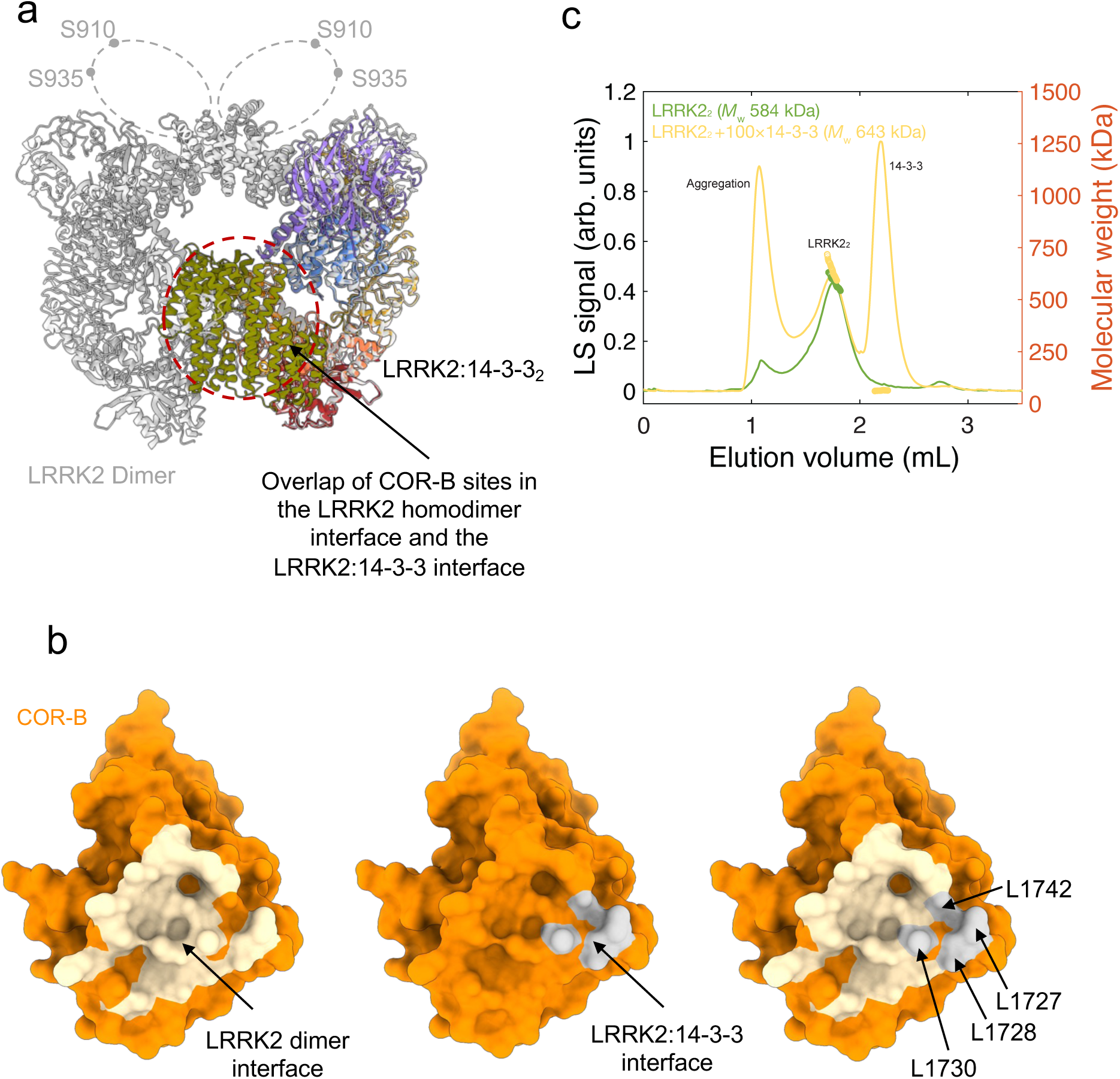
Overlap of the LRRK2:14-3-3_2_ and LRRK2 homodimer interfaces. **a.** Superimposed structures of the LRRK2:14-3-3_2_ (colored as in Fig. 1) and the dimeric structure of inactive LRRK2 (PDB: 7LHW, in gray) showing overlap of the COR-B domain mediated interfaces. The loops containing S910 and S935 sites, flexible and missing in the dimer structure, are illustrated with dashed lines. **b**. Surface representations of the COR-B domain. Left: LRRK2 dimer interface residues shaded in tan. Middle: LRRK2:14-3-3_2_ interface residues in gray. Right: Overlay showing steric interference between LRRK2 dimerization and 14-3-3 binding. **c.** SEC-MALS chromatogram for the LRRK2 dimer in the presence and absence of 14-3-3 indicating changes in molecular weight distribution.

### 14-3-3 dimer reinforces inhibition of LRRK2 kinase activity by locking its inactive conformation

In our LRRK2:14-3-3_2_ structure, the kinase domain adopts a canonical kinase-inactive conformation (Fig. 4a). This is consistent with previously reported inactive LRRK2 structures in the absence of 14-3-3^45,50,89^, where the Roc-COR domain is rotated away from the kinase domain, stabilizing it in an inactive conformation (Supplementary Fig. 7a). These observations prompted a reassessment of the role of the 14-3-3 dimer in modulating LRRK2 activity. Our structure suggests that rather than inducing an inactive LRRK2 conformation, the 14-3-3 dimer maintains this state by stabilizing the position of the LRR domain, which in turn reinforces the inactive conformation of LRRK2. The LRR domain, a critical regulatory element of LRRK2, has at least two distinct conformations corresponding to the protein’s active and inactive kinase states. In the inactive state (Fig. 1 and Fig. 4b), the LRR domain folds over and forms contacts with the kinase domain, thereby blocking substrate access and stabilizing the kinase domain in an inactive configuration. In contrast, the LRR domain dramatically repositions away from the kinase domain in the active state, as seen in a Type I inhibitor-bound form^89^ and a tetramer formed on cryo-EM grids^50^. In these active structures, both the LRR domain and the preceding loop (residues 907-982) are dynamic and absent in the density maps. (Fig. 4b). In the LRRK2:14-3-3_2_ complex, the LRR domain and the 14-3-3 dimer are situated on opposite sides of the Roc-COR-Kinase-WD40 region. The primary and secondary interactions between the loop immediately preceding the LRR domain, which contains the phosphorylation sites pS910 and pS935 for 14-3-3 binding, along with the engagement of the COR domain with 14-3-3, secures the LRR domain in an inhibitory position (Fig. 4b). This configuration prevents the LRR domain from shifting away from the kinase domain.

**Fig. 4.**
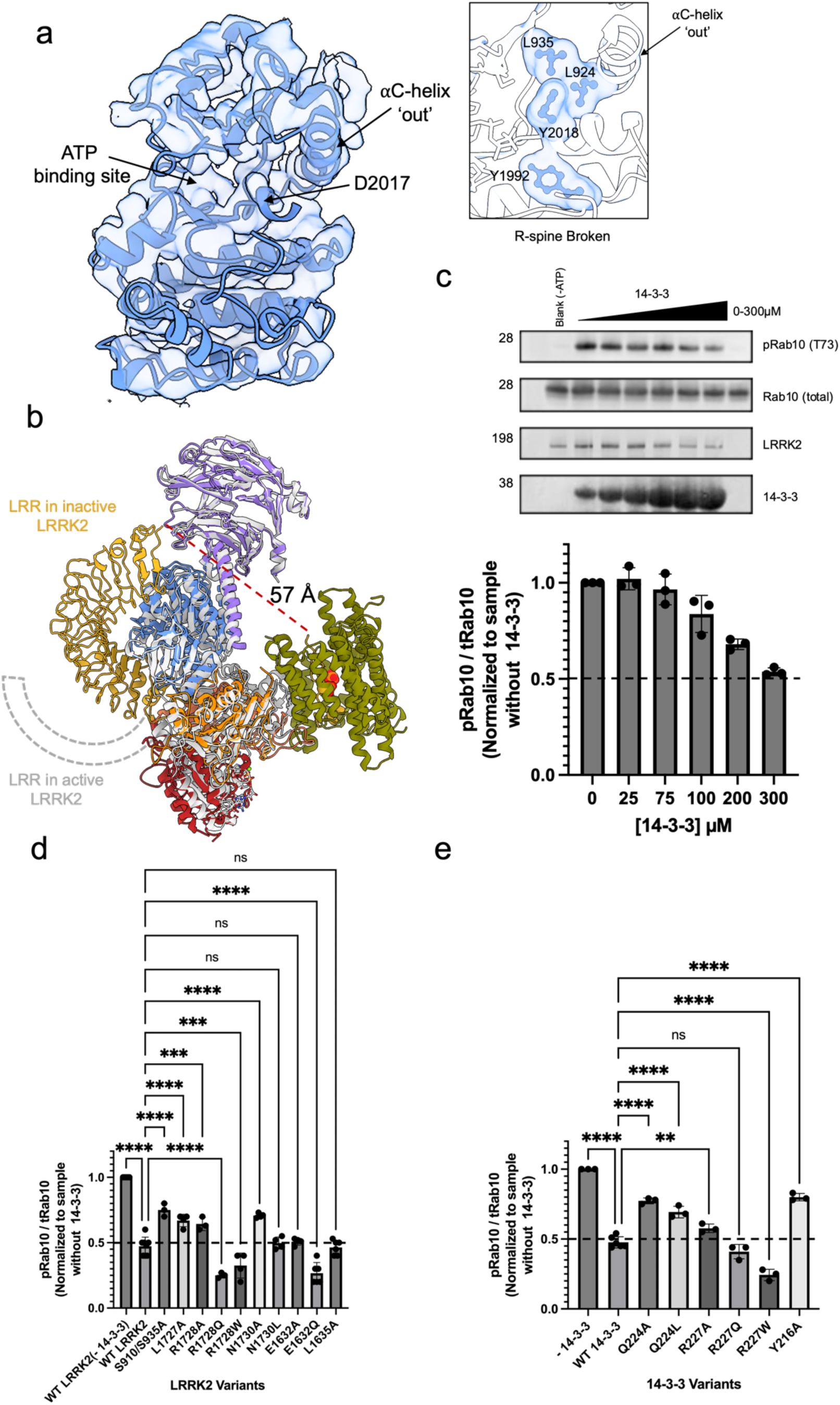
14-3-3 binding maintains LRRK2 in an inactive conformation and inhibits kinase activity. **a.** Structural representation of the kinase domain in the LRRK2:14-3-3_2_ complex showing an inactive conformation with the αC helix outward in an ‘out’ conformation and a broken regulatory (R-) spine (inset). **b.** Overlay of the LRRK2:14-3- 3_2_ complex (colored as in Fig. 1) with the active LRRK2 monomer from a LRRK2 tetramer (PDB: 8FO9, in gray), highlighting the conformational differences in the LRR domain, depicted as a cartoon. **c.** Kinase inhibition assay showing the effect of increasing concentrations of 14-3-3 on LRRK2 activity, monitored by Rab10 phosphorylation levels in a western blot. **d-e**. Impact of mutations in the LRRK2:14-3-3_2_ interface on kinase activity inhibition by 14-3-3, with results for LRRK2 mutations shown in **(d)** and 14-3-3 mutations in **(e)**. Data are mean ± SEM quantified from at least three independent protein preparations, with representative blots for inhibition assays shown in **(c).** See source data for membrane images and for significance of difference with the one-way Brown-Forsythe and Welch ANOVA test when applicable.

Additionally, we used 3D variability analysis (3DVA) in *Cryosparc*^95^ which showed that the LRR domain and the 14-3-3 are flexible relative to the Roc-COR-Kinase-WD40 region, displaying synchronized movements that approach or distance themselves from the Roc-COR-Kinase-WD40 segment. These movements restrict the motion range of the LRR domain and lock it in a kinase-inactive position (Supplementary Fig. 12, Supplementary Movie 1). This analysis demonstrates that the primary and secondary interactions between LRRK2 and 14-3-3 synergistically contribute to the inhibitory effect of 14-3-3 on LRRK2. The primary interaction recruits the 14-3-3 dimer, while the secondary interaction further stabilizes the LRR domain in a kinase-inactive configuration. To further validate the inhibitory effect of 14-3-3 on LRRK2 kinase activity, we measured Rab10 phosphorylation by purified LRRK2 using an *in vitro* kinase assay. The addition of 14-3-3 to LRRK2 inhibited Rab10 phosphorylation by up to ∼50% in a concentration-dependent manner (Fig. 4c).

We further examined the impact of mutations at the LRRK2 and 14-3-3 interfaces on LRRK2 inhibition by 14-3-3 (Fig. 4d-e). Mutants at the primary interaction residues S910/S935 and the secondary interaction residues L1727, L1828, and N1730 produced a significant reduction in LRRK2 inhibition by 14-3-3 (∼25% inhibition of Rab10 phosphorylation by 14-3-3 compared to a 50% inhibition with the wild-type (WT) protein, Fig. 4d). Similarly, mutations on the 14-3-3 interface residues Q224 and Y216 residues displayed reduced LRRK2 inhibition (∼22% inhibition of Rab10 phosphorylation by 14-3- 3 compared to a 50% inhibition with the WT protein, Fig. 4e), highlighting the importance of these interfaces in the 14-3-3 inhibitory effect. Additionally, we showed that maintaining the hydrophobic nature of the secondary interface was crucial for the interaction. When residues such as R1728 and E1632 in the COR domain and R227 in 14-3-3 were mutated to more hydrophobic ones, the inhibition was reinforced. Moreover, mutations at E1632, L1635, and S1636 in LRRK2 were not completely tolerated, as they resulted in lower expression yields or reduced thermostability (Supplementary Fig. 13 and 14).

Furthermore, we modeled the interaction of 14-3-3 with the active conformation of LRRK2 by superimposing the active conformation onto the inactive LRRK2 in the LRRK2:14-3-3_2_ complex (Fig. 5). The active LRRK2 conformation requires the Roc-COR domain to move towards the kinase domain to stabilize its active form (Fig. 5a)^50^. In this configuration, the COR-B subdomain rotates relative to the COR-A domain, making it incompatible to engage 14-3-3 simultaneously with the COR-A subdomain, as observed in the inactive conformation (Fig. 5b). This orientation prevents the complete binding of 14-3-3 to the active form of LRRK2. The widely reported loss of LRRK2 phosphorylation at S910/S935 in the presence of type I inhibitors supports this observation^61^. Type I LRRK2 inhibitors are known to stabilize LRRK2 in an active conformation^61^, albeit not able to phosphorylate downstream substrates due to the presence of ATP-competitive inhibition. However, the precise mechanisms by which these inhibitors alter LRRK2 phosphorylation and influence the stabilization of the active conformation were not well understood. Our findings suggest that *in vivo*, the destabilization of the inactive conformation and formation of an active conformation that is inhibited disrupts the LRRK2:14-3-3_2_ complex, leading to the exposure of the S910/S935 phosphorylation sites for phosphatase activity. This analysis not only elucidates the paradoxical effects of Type I inhibitors: on the one hand, they inhibit LRRK2 kinase activity by competing with ATP; on the other, they induce an active LRRK2 conformation that disrupts 14-3-3 interactions, leading to S910/S935 dephosphorylation. Additionally, it sheds light on the crucial conformational transitions between 14-3-3-bound inactive and 14-3-3-unbound active LRRK2, which are essential for understanding LRRK2 activation.

**Fig. 5.**
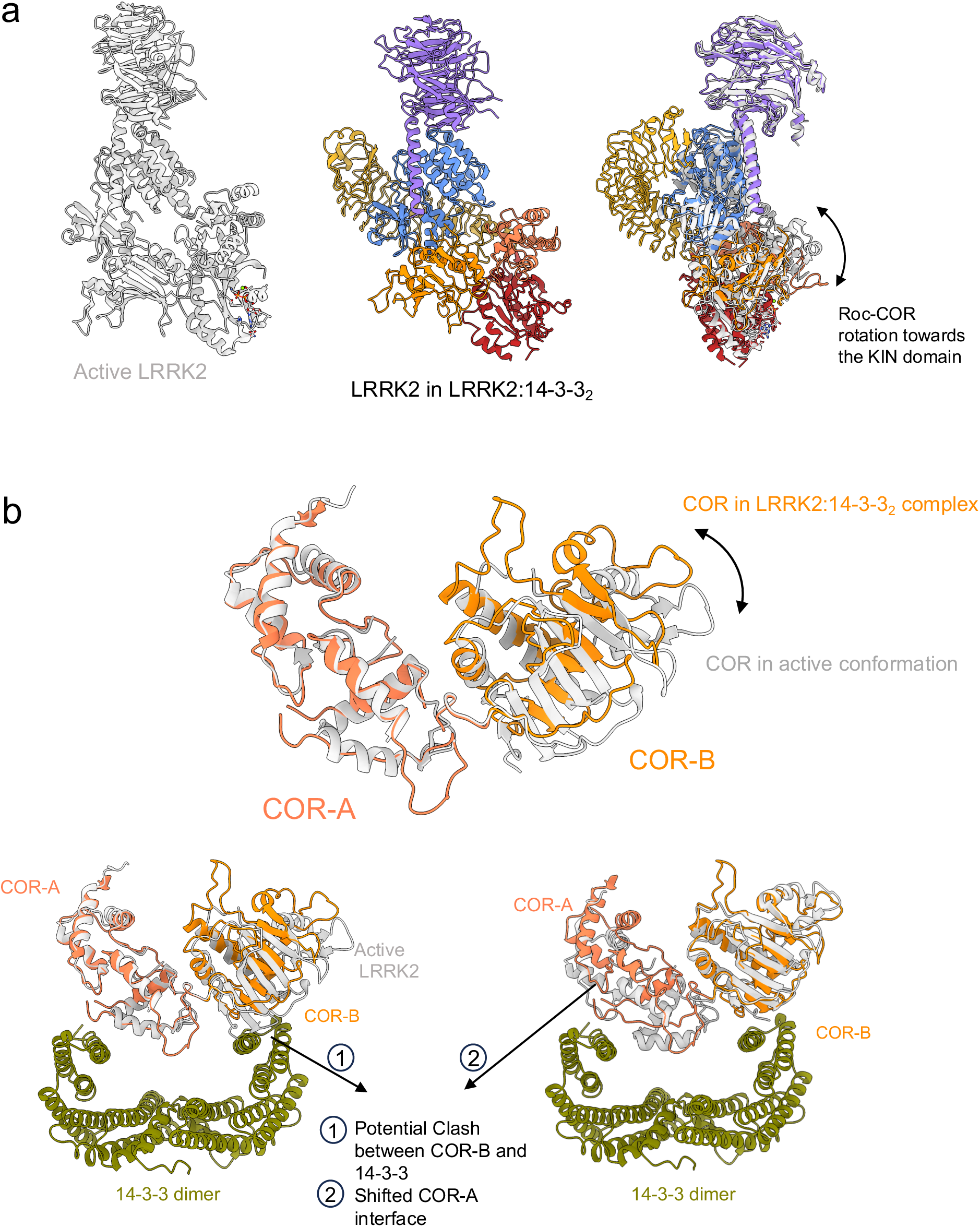
Structural comparison of the LRRK2:14-3-3_2_ complex with active LRRK2. **a.** Side by side comparison of the LRRK2:14-3-3_2_ complex (colored as in Fig. 1) alongside the active conformation of LRRK2 from a LRRK2 tetramer (PDB: 8FO9) shown in gray, highlighting the Roc-COR domain rotation (indicated by an arrow). 14-3-3 proteins are omitted for clarity. **b.** Superimposition of the COR domain from the LRRK2:14-3-3_2_ complex (shown in coral) with the COR domain from the active LRRK2 (shown in gray), alignment done by the COR-A domain (top and bottom left) and by the COR-B domain (bottom right), to illustrate the structural shifts and effects.

### Pathological mutations that disrupt complex stability and LRRK2 inhibition by 14- 3-3

Prominent PD mutations, such as Y1699C and R1441C/H/G, located in the Roc-COR GTPase domain of LRRK2, exhibit elevated kinase activity^23,94^. The COR domain, crucial for LRRK2’s functionality, participates in oligomerization interfaces for interactions with microtubules and dimer and tetramer interactions^42,45,50^. Our findings extend this understanding by demonstrating that the COR domain is critical not only for oligomerization but also for interaction with 14-3-3 proteins and that this can be disrupted by PD-related mutations. Human mutations R1628P and R1728H, located at the COR-A/14-3-3 and COR-B/14-3-3 interfaces respectively, show a twofold increase in kinase activity in *in vivo* assays^96–98^. Our *in vitro* kinase experiments corroborate this with a similar twofold increase in activity (Fig. 6a). Additionally, a ∼20% reduction in LRRK2 inhibition by 14-3-3 compared to WT was observed (Fig. 6b). Furthermore, our Co-IP experiments reveal that the mutation R1628P resulted in a ∼50% decrease in binding with endogenous 14-3-3 relative to WT (Fig. 6c), highlighting the critical impact of these interactions on LRRK2 kinase activity and PD pathogenesis. Conversely, PD-related mutations at the Roc-COR-Kinase-WD40 region, including I2020T, L1795F, Y1699C, R1441C/H/G, A1442P, and G2385R, though positioned away from the direct LRRK2:14-3-3_2_ interface, are reported to disrupt 14-3-3 interactions^23,94^. These residues are not solvent exposed and are implicated in various intramolecular interactions within the LRRK2 inactive conformation (Supplementary Fig. 10b). We explored the effects of mutations R1441G at the Roc domain and G2019S at the kinase domain on LRRK2 and 14-3-3 interaction. Both mutations exhibited approximately a 2.5-fold increase in kinase activity; strikingly, only the R1441G mutation led to a loss of interaction with 14-3-3 and a roughly 30% reduction in 14-3-3 inhibition of LRRK2 (Fig. 6), indicating that R1441G destabilizes the Roc-COR interaction in a conformation unfavorable for 14-3-3 binding. These results suggest that pathological PD mutations at the COR:14-3-3 interfaces and other parts of the Roc-COR GTPase domain disrupt 14-3-3 binding. This disruption reduces the stability of the LRRK2:14-3-3_2_ complex, leading to enhanced kinase activity, and underscores the critical role of 14-3-3 in LRRK2 regulation. Furthermore, mutations with diminished 14-3-3 interaction, such as R1441 C/G, Y1699C, and I2020T, have shown depleted levels of phosphorylation at S910/S935 sites^72,99^, possibly due to weakened secondary interactions in LRRK2:14-3-3_2_, exposing primary phospho-Ser sites for dephosphorylation. This mechanism likely contributes to the hyperactivation observed in mutations like R1628P and R1441G (Fig. 6a). Conversely, the G2019S mutation in the kinase domain, which does not significantly alter the inactive conformation of LRRK2 but most likely affects kinase kinetics^45,100^, does not affect 14-3- 3 binding (Fig. 6b).

**Fig. 6.**
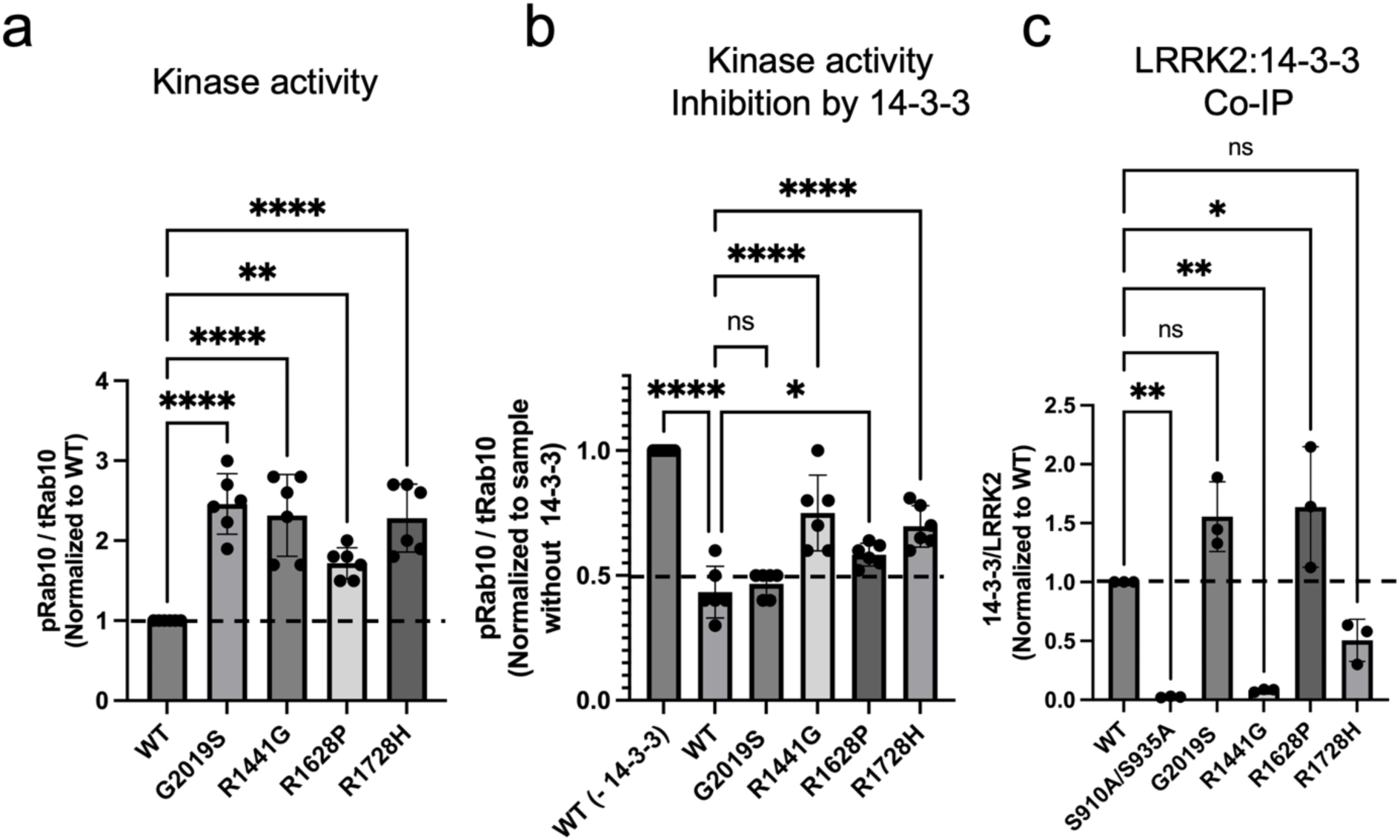
Impact of PD-associated mutations in Kinase and GTPase domains on LRRK2:14-3-3_2_ interaction. **a.** LRRK2 kinase activity assay for wild-type (WT), as a control, and various PD-related hyperactive mutants in the kinase and GTPase domains, showing increased Rab10 phosphorylation levels indicative of enhanced kinase activity. **b.** Comparative analysis of 14-3-3 inhibition of kinase activity across WT and PD mutants, illustrating that mutations result in differential effects on LRRK2 function**. c.** Co-IP assays quantifying cellular interaction between LRRK2 WT/ PD- related mutations and endogenous 14-3-3, highlighting alterations in binding affinity due to mutations. LRRK2 kinase activity data are mean ± SEM quantified from at least three independent protein preparations. Refer to Supplementary Fig. 11 and source data for membrane images and for significance of difference with the one-way Brown-Forsythe and Welch ANOVA test when applicable.

Our results indicate that not all the PD-activating mutations have the same mechanism of hyperactivation. While some mutations, like G2019S, do not significantly alter the inactive conformation of LRRK2, others like R1441G, might destabilize the overall LRRK2 inactive conformation or its dynamics, indirectly affecting the LRRK2/14-3-3 interaction and leading to LRRK2 dephosphorylation. This mechanism may be similar to the activation of Type I inhibitors, though this hypothesis merits further studies.

## Discussion

Enhanced pathological LRRK2 kinase activity is critically implicated in PD pathogenesis, highlighting the physiological need for the precise regulation of kinase activity^23,94^. Here, we elucidated the structural and mechanistic underpinning of the interactions between LRRK2 and 14-3-3 proteins. Our findings reveal 14-3-3’s dual role as an inhibitor and regulator of LRRK2, crucial for modulating LRRK2 activity under both physiological and pathological conditions.

Our structural analysis demonstrated extensive interactions between 14-3-3 proteins with key phosphorylation sites (pS910 and pS935) and the COR-A and COR-B subdomains of LRRK2 within the LRRK2:14-3-3_2_ complex. These interactions stabilize LRRK2 in an inactive monomeric conformation, restricting the mobility of the LRR region and inhibiting substrate access. In contrast in the active conformation of LRRK2, the LRR domain is repositioned to expose the kinase domain, facilitating substrate access and highlighting potential targets for interventions to maintain LRRK2 in its inactive state. Interestingly, the positioning of the LRR domain distinguishes LRRK2 from LRRK1, the other LRRK in humans^101^. Unlike LRRK2, LRRK1 lacks the N-terminus phospho-serine binding motifs necessary for the primary interaction with 14-3-3^101,102^.

Additionally, LRRK1 differs at the secondary interface, which helps explain why LRRK1 is not regulated by 14-3-3 in the same way as LRRK2 (Supplementary Fig. 10c). This underscores a fundamental divergence in the regulatory mechanisms of LRRK1 and LRRK2.

Our mutagenesis studies, both *in vitro* and in cells, revealed that pathological mutations at the COR:14-3-3 interfaces and other areas within the Roc-COR GTPase domain substantially reduce 14-3-3 binding and subsequently enhance LRRK2 kinase activity. This emphasizes the crucial role of 14-3-3 interactions in controlling LRRK2 activity. Our results also indicate that 14-3-3 binding maintains LRRK2 in its inactive conformation and prevents the formation of LRRK2 dimers and higher oligomers. This suggests that the LRRK2:14-3-3_2_ complex likely represents LRRK2’s dormant state within cells, given the relatively high endogenous expression levels of 14-3-3 compared to LRRK2.

Furthermore, our analyses indicate that the COR conformation in the active state of LRRK2 prevents simultaneous engagement with both COR-A and COR-B, blocking stable complex formation with active LRRK2. This is supported by the observed reduction in LRRK2 phosphorylation at S910/S935 in the presence of Type I inhibitors^61^, which favor the active conformation of LRRK2. This suggests that the active conformation promoted by these inhibitors likely disrupts both secondary and primary interactions with 14-3-3 proteins, potentially altering LRRK2’s functional state and impacting its regulatory mechanisms.

Given these complexities, critical questions about LRRK2’s functional states arise: How is LRRK2 converted from its 14-3-3 bound inactive form to an active form, and what constitutes LRRK2’s physiological active form? It is possible that the monomeric, active form of LRRK2, which is free from 14-3-3 binding but may interact with other cofactors or be favored under specific cellular conditions, represents its most active state. Further experimental exploration is necessary to fully understand LRRK2 dynamics and activation mechanisms.

Our results have significant implications for PD biomarkers and therapeutics. Phosphorylation levels of LRRK2 at S910/S935 and LRRK2 substrates Rab8/10 are used as biomarkers in clinical trials for monitoring PD progression and therapeutic responses^14,103–107^. Studies have shown that dephosphorylation at S910/S935 and elevated Rab8/10 phosphorylation occurs in both familial and sporadic PD^40^. Here, we provide mechanistic insights into how phosphorylation at these sites is important for LRRK2’s interaction with 14-3-3 proteins to maintain its inactive state and how the active conformation of LRRK2 can promote the exposure of these sites for dephosphorylation, leading to elevated Rab substrate phosphorylation levels.

We further explain how Type I inhibitors reduce S910/S935 phosphorylation levels, which have been used to measure target engagement and therapeutic response in trials of LRRK2 inhibitors. These inhibitors stabilize LRRK2 in an active conformation that is ATP-competitively inhibited, thus impairing its 14-3-3 binding and making pS910/pS935 sites more susceptible to dephosphorylation. Based on our findings, we proposed that the use of S910/S935 phosphorylation levels may need to be tailored to specific LRRK2 mutations and idiopathic PD conditions that promote the active conformation of LRRK2 and impair 14-3-3 binding. Additionally, the use of S910/S935 phosphorylation levels as biomarkers may need to be tailored to assess therapeutic responses to inhibitors that trap LRRK2 in an active conformation, thereby reducing 14-3-3 binding. Our results provide a structural basis for understanding key aspects of PD biomarkers, offering a targeted approach to patient stratification in clinical settings.

Based on our findings, we also propose developing LRRK2 and 14-3-3 protein-protein interaction stabilizers, termed ’glue’ molecules^108–110^, specifically designed to enhance the interaction between LRRK2 and 14-3-3. Such molecules could stabilize the autoinhibitory complex, thereby reducing the pathological activity of LRRK2 mutations and presenting a novel therapeutic strategy for mitigating PD’s effects.

In summary, our study establishes a comprehensive structural framework for understanding how 14-3-3 proteins regulate LRRK2 and how PD associated mutations impact their interactions, paving the way for new therapeutic strategies targeting this crucial interaction. Given LRRK2’s significant role in neurodegenerative disorders, our work highlights the key aspects of PD biomarkers and the therapeutic potential of modulating protein-protein interactions and offers a promising strategy to mitigate the detrimental effects of LRRK2 in PD and related diseases.

## Methods

### Reagents and resources

Resources used in this study are listed in Supplementary Table 1. Plasmids and cell lines are available for use upon reasonable request to the corresponding author.

### Cell lines and culture conditions

Expi293F^TM^ cells (Thermo Fisher Scientific, cat. A14527) were cultured in Expi293^TM^ and FreeStyle^TM^ 293 expression medium and cultured at 37°C with 8% CO_2_.

### Expression and purification of LRRK2

Full-length human LRRK2 (Uniprot Q5S007 with R85H mutation), with an N-terminus 3X-FLAG and a rhinovirus 3C protease cleavage site, was codon-optimized for *Homo Sapiens* and synthesized by GenScript. The gene was sub-cloned into the pEG BacMam vector for mammalian cell protein expression^111^. Bacmid was generated using the Bac-to-Bac system from Invitrogen and transfected into Sf9 cells for baculovirus generation. Expi293F cells were seeded at 2.0 x 10^6^ cells/mL in four liters of FreeStyle^TM^ 293 expression medium and infected with high titer baculovirus. After ∼12 hours of incubation at 37°C with shaking, 10mM sodium butyrate was added and the temperature was reduced to 30°C. Cells were collected by centrifugation after 72 hours of incubation, and cell pellets were flash frozen for later purification or resuspended in resuspension buffer (20 mM Tris pH 8.3, 10 mM CaCl_2_, 5 mM MgCl_2_, 100 mM NH4Cl,

100 mM NaCl, 10 mM β-glycerophosphate and 1 mM sodium vanadate 50 mM L-Arg, 50 mM L-Glu, 0.0008% Tween-80, 10% glycerol). All subsequent purification steps were carried out at 4°C. Cells were lysed using sonication, and the clarified lysate was incubated with FLAG-M2 affinity resin (Sigma) for 2 hours with rotation. The resin was washed once with resuspension buffer, twice with wash buffer (20 mM Tris pH 7.4, 500 mM NaCl, 5 mM MgCl2, 0.0008% Tween-80), and then once with gel filtration buffer (20 mM HEPES pH 8.3, 150 mM NaCl, 5 mM MgCl2, 0.0008% Tween-80). For elution, the column was washed three times with gel filtration buffer supplemented with 150 μg/mL 3X-FLAG peptide. The eluted protein was concentrated to ∼500 µL and injected onto a Superose 6 Increase 10/30 column (Cytiva) equilibrated in gel filtration buffer. Peak fractions corresponding to LRRK2 monomer were collected for complex formation with 14-3-3 protein. LRRK2 mutants were obtained from GenScript as well and affinity purified as described for WT LRRK2. Protein concentrations were determined using extinction coefficients calculated from the protein sequence.

For kinase assays WT and LRRK2 variants were expressed in a small-scale format with 50-200 mL cell cultures, protein expression, and purification were carried out as described above with the following changes. Lysis was carried out by incubating in resuspension buffer supplemented with 1% Tween-80 at 4 °C for 1 hour. After clarification, the lysate was incubated with 50 μL FLAG-M2 affinity resin for 2 hours, followed by washes as described above. A single protein elution was done with gel filtration buffer supplemented with 150 μg/mL 3X-FLAG peptide. The eluted protein was filtered through a 0.22 μm Ultrafree-GV centrifugal filter (Sigma cat. UFC30GVNB) to remove residual resin before use in kinase and inhibition assays.

### Expression and purification of 14-3-3γ

N-terminus His tagged 14-3-3 γ (Uniprot P61981), with a tobacco etch virus (TEV) protease cleavage site, codon-optimized for *Escherichia coli*, was synthesized by Genscript and subcloned into a pET21b vector. This construct was transformed into *E. coli* BL21*(DE3) cells for protein expression. The starter cultures were grown in LB medium with 0.1 mg/ml ampicillin overnight at 37 °C and then 1:100 diluted into the same medium. The cultures were grown at 37° C until the OD_600_ reached ∼0.5-0.6.

Protein expression was induced by adding 1mM isopropyl β-D-thiogalactoside (IPTG) and the cultures were subsequently incubated for 18-20 hours at 16 °C. Cells were harvested by centrifugation, resuspended in lysis buffer (25 mM Tris-HCl pH 7.4, 150 mM NaCl, 20 mM Imidazole, 1 mM TCEP, and protease inhibitors from Roche), and lysed by several passes through a microfluidizer (LM20 Microfluidizer, Microfluidics Corp). All subsequent purification steps described below were carried out at 4°C. The lysates were centrifuged, and the collected supernatants were incubated with Ni-nitrilotriacetic (Ni-NTA) agarose beads for 0.5 hours. The beads were washed with five column volumes of lysis buffer. The protein was eluted with elution buffer (25 mM Tris-HCl pH 7.4, 150 mM NaCl, 500 mM Imidazole, 1 mM TCEP) in a gradient over twenty column volumes. Fractions containing 14-3-3 were pooled, concentrated to ∼5mL, and injected onto a Superdex 75 16/160 column (GE Healthcare) which was equilibrated in gel filtration buffer. Fractions containing purified 14-3-3 were pooled, concentrated to ∼10 mg/mL, and stored at −80 °C for long-term use. Mutants of 14-3-3 were generated using the NEB Q5 Site-Directed Mutagenesis Kit (cat. E0554). Protein concentrations were determined using UV absorbance at 280 nm, using the extinction coefficients calculated from the protein sequence.

### Expression and purification of Rab10

N-terminus His-SUMO tagged Rab10 (Uniprot P61026), codon-optimized for expression in *E. coli*, was synthesized by Genscript and subcloned into a pET21b vector. The constructs were transformed into *E. coli* BL21 cells. Starter cultures were grown in LB medium supplemented with 0.1 mg/ml Ampicillin at 37 °C overnight and then 1:100 diluted into the same medium. The cultures were grown at 37° C until the cell density reached ∼0.5-0.6 OD_600_ and induced with 1mM isopropyl β-D-thiogalactoside (IPTG) and continued at 19 °C for 18-20 hours. Cells were harvested by centrifugation, resuspended in lysis buffer (25mM Tris 8.0, 500mM NaCl, 1mM MgCl2, 1mM TCEP, and protease inhibitors from Roche), and lysed by sonication. All subsequent purification steps described below were carried out at 4°C. The lysates were centrifuged, and the collected supernatants were incubated with Ni-nitrilotriacetic (Ni-NTA) agarose beads for 0.5 hours.

The beads were washed with 60 column volumes of lysis buffer, followed by 20 column volumes each of lysis buffer supplemented with 10 mM and 25 mM imidazole, respectively. The protein was eluted with elution buffer (25 mM Tris-HCl pH 8.0, 100 mM NaCl, 500 mM Imidazole, 1 mM TCEP) over a gradient of twenty column volumes.

Eluates were spin dialyzed into the lysis buffer, after which NP-40 was added to a final concentration of 0.1% and subjected to UlP1 (an engineered SUMO protease) digestion for 1 hour at 25 °C at a molar ratio of 1: 200 (protein:enzyme) to cleave the His-SUMO tag. The cleaved tag and the protease were then removed using Ni-NTA beads. The Rab10 fractions were pooled, concentrated to ∼5mL, and injected onto a Superdex 75 16/160 column (GE Healthcare) equilibrated in gel filtration buffer. Fractions containing purified Rab10 were taken, pooled, concentrated to ∼10 mg/mL, and stored at −80 °C for long-term use. Protein concentrations were determined using UV absorbance at 280 nm, using the extinction coefficients calculated from the protein sequence.

### LRRK2/14-3-3 complex formation

Purified LRRK2 monomer fraction was diluted in gel filtration (GF) buffer with no NaCl to reduce NaCl concentration to 50mM, and then incubated with at least 60-fold molar excess of 14-3-3 protein on ice for 15 minutes to promote complex formation. For experiments requiring cross-linking, the protein mixture was first incubated for 10 minutes at room temperature, followed by the addition of 1 mM bis(sulfosuccinimidyl)suberate (BS3). After a further 15-minute incubation at room temperature, the reaction was quenched with 50mM Tris, pH 7.4. The cross-linked LRRK2/14-3-3 complex was concentrated to ∼500 µL and injected onto a Superose 6 Increase 10/30 column (Cytiva) which was equilibrated in gel filtration buffer. Fractions from the peak corresponding to the cross-linked LRRK2/14-3-3 complex were used for cryo-EM grid preparation.

### Mass photometry

Samples containing LRRK2 or LRRK2/14-3-3 complexes were diluted to concentrations ranging from 20-50nM in detergent-free gel filtration buffer for mass photometry measurements using a OneMP mass photometer (Refeyn). Movies were collected for 6,000 frames over 60 seconds in regular view mode. Mass determination was conducted using the DiscoverMP software, with calibration performed using a mixture of beta amylase (Sigma, cat. A8781) and thyroglobulin (Sigma, cat. T9145).

### Multi-angle static light scattering (MALS)

SEC-MALS data were collected using a Shimadzu LC-20AD HPLC, coupled to a Shimadzu SPD-20A UV detector, a Wyatt Dawn MALS detector, and a Wyatt Optilab refractive index detector. Data were collected following in-line fractionation with a Superose 6 Increase 15/150 column (GE Healthcare), pre-equilibrated in gel filtration buffer, operated at a flow rate of 0.3 mL/min. 50 μL of the dimeric LRRK2 at 130 nM were applied to the column for analysis in the absence and presence of increasing concentrations of 14-3-3γ ranging from 1-100 times that of LRRK2. Data were processed using ASTRA software v. 8.0.2.5 (Wyatt). The detector response was normalized using monomeric BSA (Thermo Fisher, cat. 23209). Protein concentration was determined using differential refractive index, using a dn/dc value of 0.185 mL/g.

### LRRK2 kinase activity assays and inhibition assays with 14-3-3

Kinase reaction mixtures consisted of 100 nM LRRK2 and 3 µM Rab10, in 20 mM HEPES buffer pH 7.4, 50 mM NaCl, 5 mM MgCl_2_, and 0.0008% Tween-80. Reactions were carried out at 30 °C for 30 minutes in a thermomixer (Eppendorf) with shaking (300 rpm). The reactions were initiated by adding 5mM ATP and terminated by the addition of LDS-NuPAGE loading buffer (Thermofisher). The samples were boiled at 95 °C for 10 minutes and stored at -80 °C if not processed immediately in immunoblot analysis. Kinase assays were performed using at least three independent protein preparations, each in duplicate. The samples were resolved on 4-12% bis-tris gels and wet-transferred to a 0.4 µm PVDF membrane. Membrane blocking was done with 5% bovine serum albumin (BSA) in TBS-T (20 mM Tris, 150 mM NaCl pH 7.4, 1% Tween- 20) for 30 minutes at room temperature before probing with anti-phospho Rab10 T73 primary antibody (Abcam, cat. ab241060, dilution 1:500), anti-Rab10 primary antibody (Abcam cat. ab237703, dilution 1:1000), and anti-LRRK2 primary antibody (Abcam cat. ab133474, dilution 1:25000). Membranes were washed with TBS-T three times and then incubated with goat anti-rabbit and anti-mouse IR-fluorescent secondary antibodies (Li-Cor, cat. 926-3221, Li-Cor, cat. 926-68072) for 1 hour at room temperature. Following four washes, membranes were imaged using a Typhoon scanner (GE Healthcare, software v. 1.1.0.7). Blots were quantified using ImageStudio Lite software (v.5.2.5) to determine the pRab10/Rab10 ratio, normalized to WT controls on the same membrane. Statistical significance was quantified using a one-way Brown-Forsythe and Welch ANOVA test in Prism (v.10.2.0), with pairwise comparisons via unpaired t-tests with Welch’s correction.

For inhibition assays with 14-3-3 proteins, kinase reactions were carried out as described above with an additional incubation of the reaction mixture with 14-3-3 on ice for 15 minutes prior to ATP addition. The dependency of LRRK2 inhibition on 14-3-3 concentration was evaluated in a range from 0 to 300 µM. The effects of mutations in LRRK2 or 143-3-3 on inhibition were tested using 14-3-3 at 300µM. Additionally, after Rab10 phosphorylation probing, blots were further analyzed with anti-14-3-3 gamma antibody (Abcam, cat. ab137048, dilution 1:1000).

### Cryo-EM grid preparation, data acquisition, and processing

After gel filtration, the LRRK2/14-3-3 complexes were concentrated to ∼0.15-0.20 mg/mL using a 100K pore size Pall Microsep™ advance centrifugal device. Quantifoil Au R1.2/1.3 holey carbon grids, 300 mesh, were glow-discharged for 30 seconds at 25 mA on both sides. 1.5 µL of protein solution was applied to each side of the grids. Grids were vitrified using a Leica EM GP2 plunge freezer with a blotting time of 1-3 seconds. Cryo-EM data acquisition was conducted at the cryo-EM facility in the Center for Structural Biology, NCI-Frederick, using a Talos Arctica G2 (Thermo Fisher) equipped with a Gatan K3 direct detector, and energy filter, operated at 200 keV. Data were collected in super-resolution mode at a nominal magnification of 100,000×, corresponding to 0.405 Å/pixel. 50 frames per movie were acquired for a total dose of approximately 50 elections/Å^2^. Data collection was managed with EPU software (Thermo Fisher), setting defocus values ranging from -0.8 to -2.5 µm. Cryo-EM data analysis was performed using Cryosparc 3.3^95^. Movies were imported, patch-motion and patch-CTF corrected. Movies were binned to the physical pixel size in the patch motion step. Micrographs with CTF resolution > 5Å or with visible bad ice were excluded. An initial subset of particles was picked using blob picker and used to train a Topaz^112,113^ model that was used to pick particles in the entire data set. Particles were curated using multiple rounds of 2D classifications, after which duplicated particles were removed. An initial 3D classification with Ab-initio and heterogeneous refinement^114^ identified distinct volumes corresponding to unbound LRRK2, LRRK2:14-3-3_2_ complex, and LRRK2 dimer. These volumes were used for multiple rounds of heterogeneous refinement. Further non-uniform refinement ^114^ of the LRRK2:14-3-3_2_ class achieved a map with 3.96 Å resolution, as determined by the gold standard FSC. The structural dynamics of the complex were analyzed using 3D variability analysis^115^ with a 7 Å filter. Local refinement of the Roc-COR:14-3-3 portion of the map improved the resolution to 3.87 Å. Graphical summary of the cryo-EM data processing is presented in Supplementary Fig. 5c. Cryo-EM reconstruction statistics are detailed in Supplementary Table 2. The global refined map was additionally post-processed using deepEMhancer v0.13^116^. The refined maps were deposited in the EMDB database. The LRRK2:14-3-3_2_ complex model was built by the rigid-body fitting of individual monomeric proteins, LRRK2 (PDB 7LHW, modified to include T1647S and T2397M mutations as in WT sequence and have residues 540 to 703 deleted) and 14-3-3γ (PDB 2B05), into the global refined map. Each protomer of the 14-3-3 dimer was fit individually, and the LRRK2 phospho-binding sites on each 14-3-3 protomer were built manually. Fitting the models into their respective maps was initially done using UCSF Chimera^117^. Manual adjustments of the model were performed in Coot^118^ using the global and local refined maps, followed by iterative rounds of real-space refinement in Phenix^119^ and manual fitting in Coot^118^. Model validation was performed based on statistics from Ramachandran plots and MolProbity scores from Phenix and Coot^120,121^. Statistics for the final refinements are presented in Supplementary Table 2. Figures were generated by UCSF ChimeraX^122^. Figures and structural analyses, including structure deviations and electrostatic potential of surfaces were generated and calculated employing MatchMaker and Coulombic Surface plugins, respectively, using UCSF ChimeraX. The final model was deposited in the PDB database.

### Differential scanning fluorometry

Gel filtration LRRK2 and LRRK2/14-3-3 complex samples, at a concentration of ∼0.1 mg/ml, were subjected to thermal stability measurements in a Prometheus NT.48 nano- DSF instrument (NanoTemper) using nanoDSF glass capillaries. Thermal unfolding of the proteins was measured by heating the samples at a rate of 1 °C/min. The protein melting temperatures were calculated by analyzing the first derivative of the ratio of tryptophan fluorescence intensities at 330 nm and 350 nm.

### Mass Spectrometry data acquisition and analysis

LRRK2:14-3-3_2_ samples were prepared by reducing with 3 mM Tris(2- carboxyethyl)phosphine (TCEP) hydrochloride at room temperature for 1 hr, followed by alkylation with 5 mM N-Ethylmaleimide for 10 min. Proteins were then digested with trypsin (Trypsin Gold, Mass Spectrometry Grade, Promega) using a 1:20 enzyme to sample ratio (w/w) at 37 °C for 18 hr. The digested samples were desalted using a µElution HLB plate (Waters). Mass spectrometry data acquisition was performed on a system where an Ultimate 3000 HPLC (Thermo Scientific) was coupled to an Orbitrap Lumos mass spectrometer (Thermo Scientific) via an Easy-Spray ion source (Thermo Scientific). Peptides were separated on an ES902 Easy-Spray column (Thermo Scientific). The composition of mobile phases A and B was 0.1% formic acid in HPLC water, and 0.1% formic acid in HPLC acetonitrile, respectively. The mobile B amount was increased from 3% to 20% in 63 minutes at a flow rate of 300 nL/min. The Thermo Scientific Orbitrap Lumos mass spectrometer was operated in data-dependent mode.

The MS1 scans were performed in orbitrap with a resolution of 120K at 200 m/z and a mass range of 375-1500 m/z. MS2 scans were conducted in ion trap. Higher energy collisional dissociation (HCD) method was used for MS2 fragmentation with normalized energy at 32%.

Database search was performed with Proteome Discoverer 2.4 software using the Mascot search engine, against a house-built database containing the sequences of interest and Sprot Human database. The mass tolerances for precursor and fragment were set to 5 ppm and 0.6 Da, respectively. Up to 2 missed cleavages were allowed for data obtained from trypsin digestion. NEM on cysteines was set as a fixed modification. Variable modifications include Oxidation (M), Met-loss (Protein N-term), Acetyl (Protein N-term), and Phosphorylation (STY). Peptides matched with phosphorylation modification were manually curated.

### Co-immunoprecipitation and western blot analysis of LRRK2 and 14-3-3

HEK293FT cells transfected with WT or mutant LRRK2 plasmids were lysed in lysis buffer (20 mM Tris-HCl pH 7.5, 150 mM NaCl, 1 mM EDTA, 0.3% Triton X-100, 10% Glycerol, 1x Halt phosphatase inhibitor cocktail from Thermo Scientific and protease inhibitor cocktail from Roche) for 30 min on ice. The lysates were centrifuged at 4°C for 10 minutes at 20.000 g, and the supernatant was further cleared by incubation with Easy view Protein G agarose beads (Sigma-Aldrich) for 30 min at 4°C. After removing the agarose beads by centrifugation, the supernatants were incubated with FLAG-M2 affinity resin (Sigma) for 1 hour at 4°C on a rotator. The beads were washed four times with wash buffer (20 mM Tris-HCl pH 7.5, 150 mM NaCl, 1 mM EDTA, 0.1% Triton X- 100, 10% Glycerol) and eluted in elution buffer (25 mM Tris-HCl, pH 7.5, 5 mM beta-glycerophosphate, 2 mM dithiothreitol DTT, 0.1 mM Na_3_VO_4_, 10 mM MgCl2, 150 mM NaCl, 0.02% Triton and 150 ng/µl of 3X-FLAG peptide (Sigma-Aldrich)) by shaking for 30 minutes at 4°C. Each co-immunoprecipitation was quantified as a ratio between two immunoprecipitated proteins, endogenous 14-3-3 and LRRK2. Proteins were resolved on 4–20% Criterion TGX pre-cast gels (Bio-Rad) in SDS/Tris-glycine running buffer and transferred to membranes using the semi-dry trans-Blot Turbo transfer system (Biorad). Membranes were blocked with Odyssey Blocking Buffer (Li-Cor cat. 927-40000) and then incubated overnight at 4°C with the primary antibodies for anti-LRRK2 (Abcam cat. ab133474, dilution 1:2000) and anti-pan-14-3-3 (Santa Cruz cat. sc-133233, dilution 1:2000) and anti-Cyclophilin B (Abcam cat. ab16045, dilution 1:2000). The membranes were washed in TBS-T three times for 5 min followed by incubation for 1h at room temperature with fluorescently conjugated goat anti-mouse or rabbit IRDye 680 or 800 antibodies (Li-Cor). The blots were washed in TBST three times for 5 min at room temperature and scanned on an ODYSSEY® CLx (Li-Cor). Quantitation of western blots was performed using Image Studio (Li-Cor).

## Data availability

The cryo-EM density maps were deposited in the Electron Microscopy Data Bank (EMDB) with the accession code EMD-45609 for the LRRK2:14-3-3_2_ global refinement map. The corresponding atomic model for the LRRK2:14-3-3_2_ complex was deposited in the Protein Data Bank (PDB) under the accession code 9CI3. Additional data generated during this study, including results from kinase assays, inhibition assays, coimmunoprecipitation, mass spectroscopy, SEC-MALS, and differential scanning fluorimetry melting temperature data, are provided in the Source Data files.

## Acknowledgments

This research was supported by Federal funds from the National Cancer Institute, National Institutes of Health, under project number ZIA BC 011744 (P.Z.), and was additionally supported in part by the Intramural Research Program of the National Institute on Aging, National Institutes of Health. This work utilized the Center for Structural Biology cryo-EM facility, NCI at Frederick, and we would like to thank Dr. Dan Shi for his assistance with cryo-EM data collection. We also thank Dr. Yan Li from the protein /peptide sequencing facility at the National Institute of Diabetes and Digestive and Kidney Diseases for mass spectrometry data collection. We thank Dr. Sergey G. Tarasov and Marzena Dyba for assistance with collecting the differential scanning fluorometry and mass photometry data in the Biophysics Resource at the Center for Structural Biology, NCI at Frederick. Additionally, this study utilized the computational resources of the Frederick Research Computing Environment cluster.

## Author Contributions

J.A.M.F. and P.Z. conceptualized the project. J.A.M.F. collected and processed the cryo-EM data with input from R.M. and P.Z. J.A.M.F, N.L. and A.A.C prepared all recombinant protein samples. J.A.M.F. undertook the biochemical/biophysical characterization of protein samples. J.A.M.F and R.M. performed the MALS experiment. A.B. performed the in vitro cell assays. M.R.C and P.Z. supervised the work. J.A.M.F. and P.Z. wrote the first draft of the manuscript, and all authors approved the final version of the manuscript.

## Declaration of Interests

The authors declare no competing interests.

## Supplementary Materials

**Supplementary Fig. 1.**
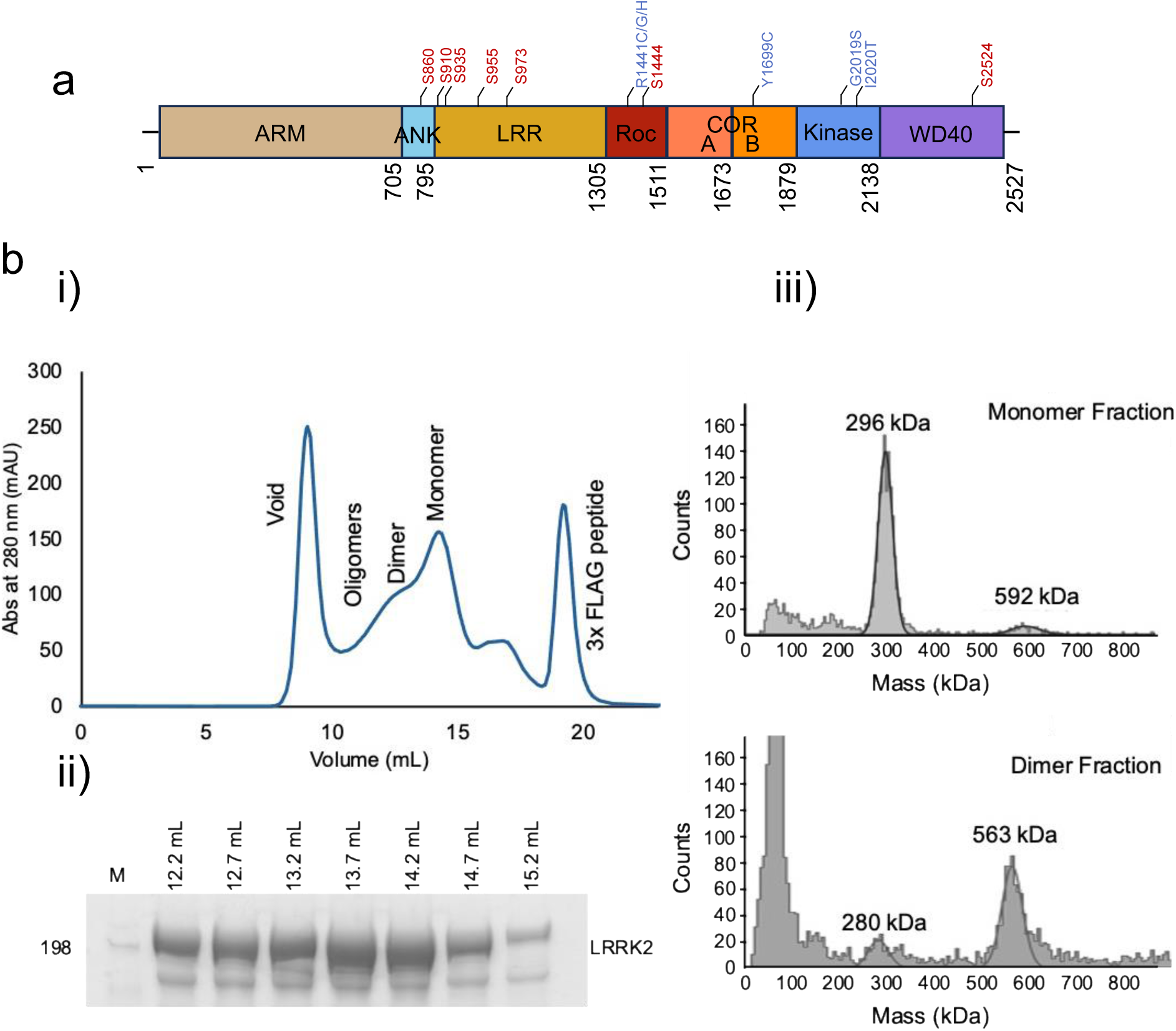
Isolation and characterization of human LRRK2. **a.** Schematic of LRRK2 domain organization with potential 14-3-3 phospho-binding sites (marked in red) and prominent PD mutations (marked in blue). **b.** Purification profile of LRRK2. (i) Elution profile from a Superose 6 Increase 10/30 gel filtration column indicating the monomeric and dimeric states of LRRK2 (ii) Corresponding SDS-PAGE gel analysis of LRRK2 fractions collected during gel filtration (iii) Mass photometry analysis of the monomer (top) and dimer (bottom) fractions.

**Supplementary Fig. 2.**
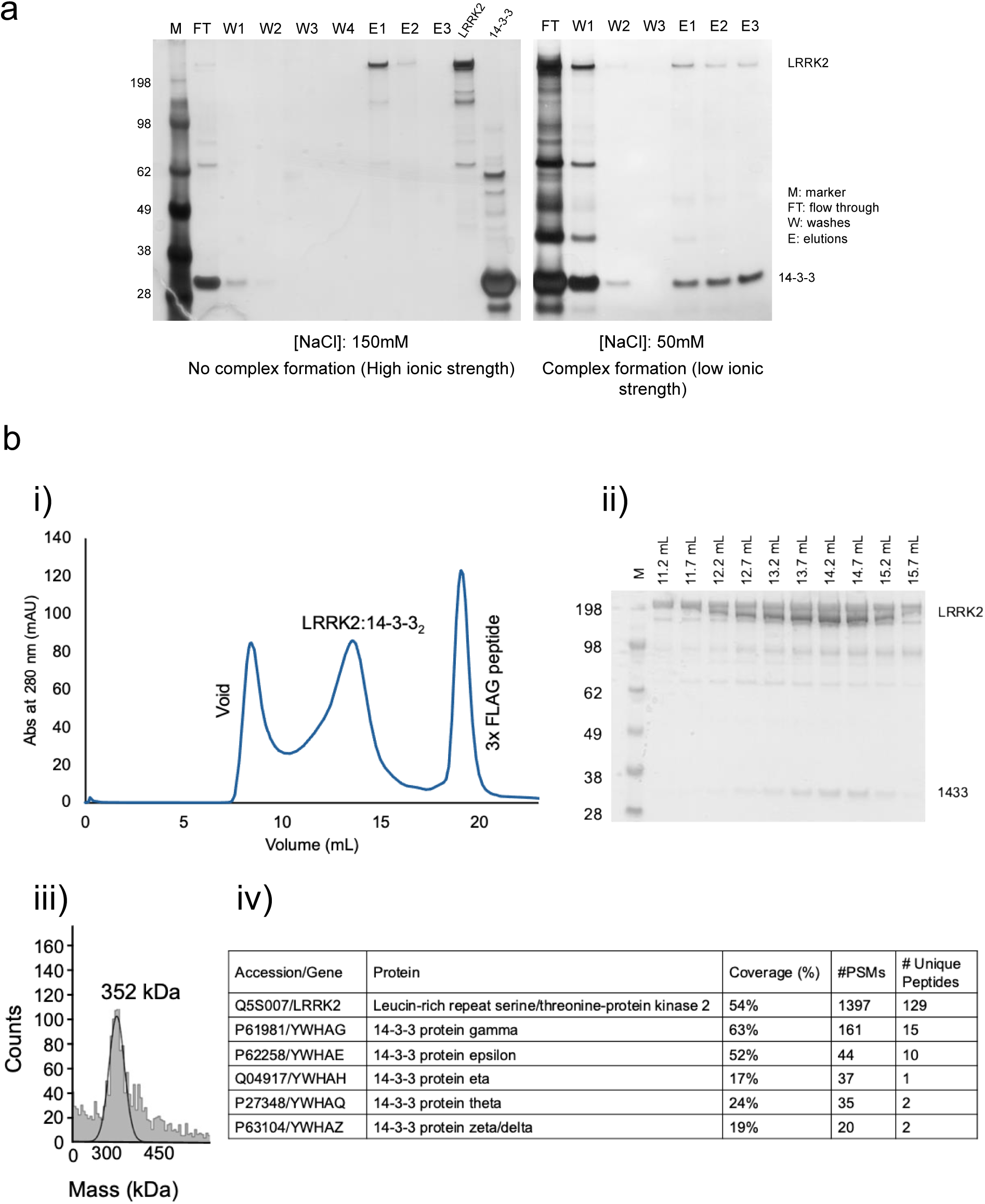
Isolation and characterization of human LRRK2/14-3-3 complex. **a.** Dependency of LRRK2/14-3-3 complex formation on the ionic strength of the solution. Co-IP assays using purified FLAG-LRRK2 to capture 14-3-3γ showed that LRRK2 does not form a stable complex with 14-3-3 at high salt concentration (left gel), while a stable complex is formed under low ionic strength conditions (right gel), illustrating the ionic sensitivity of this interaction. **b.** Biochemical characterization of the LRRK2/14-3-3 complex: (i) Elution profile from Superose 6 Increase 10/30 gel filtration column indicating the presence of the LRRK2:14-3-3_2_ complex. (ii) SDS-PAGE analysis validates the co-elution of LRRK2 and 14-3-3 proteins in the corresponding peak fraction. (iii) Mass photometry measurements confirm the stoichiometry of the complex. (iv) Mass spectrometry analysis of the complex in solution. Number of total (Peptide Spectrum Matches (PSMs)) and unique peptides are reported.

**Supplementary Fig. 3.**
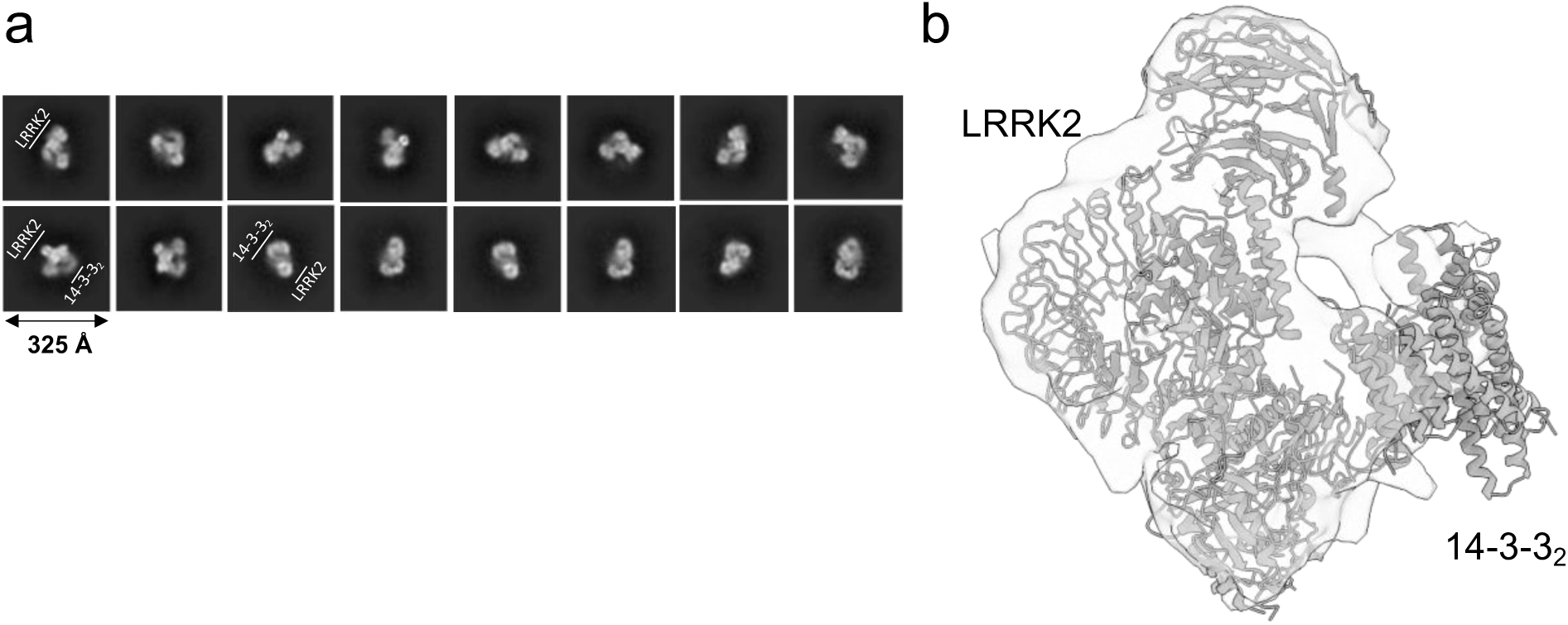
Diagnostic cryo-EM analysis of the LRRK2/14-3-3 complex. Representative 2D class averages **(a)** and cryo-EM density map at 9.4 Å resolution **(b)** of the LRRK2/14-3-3 complex, with the monomeric LRRK2 (PDB: 7LHW) and 14-3-3 dimer (PDB: 2B05) are fitted within the map.

**Supplementary Fig. 4.**
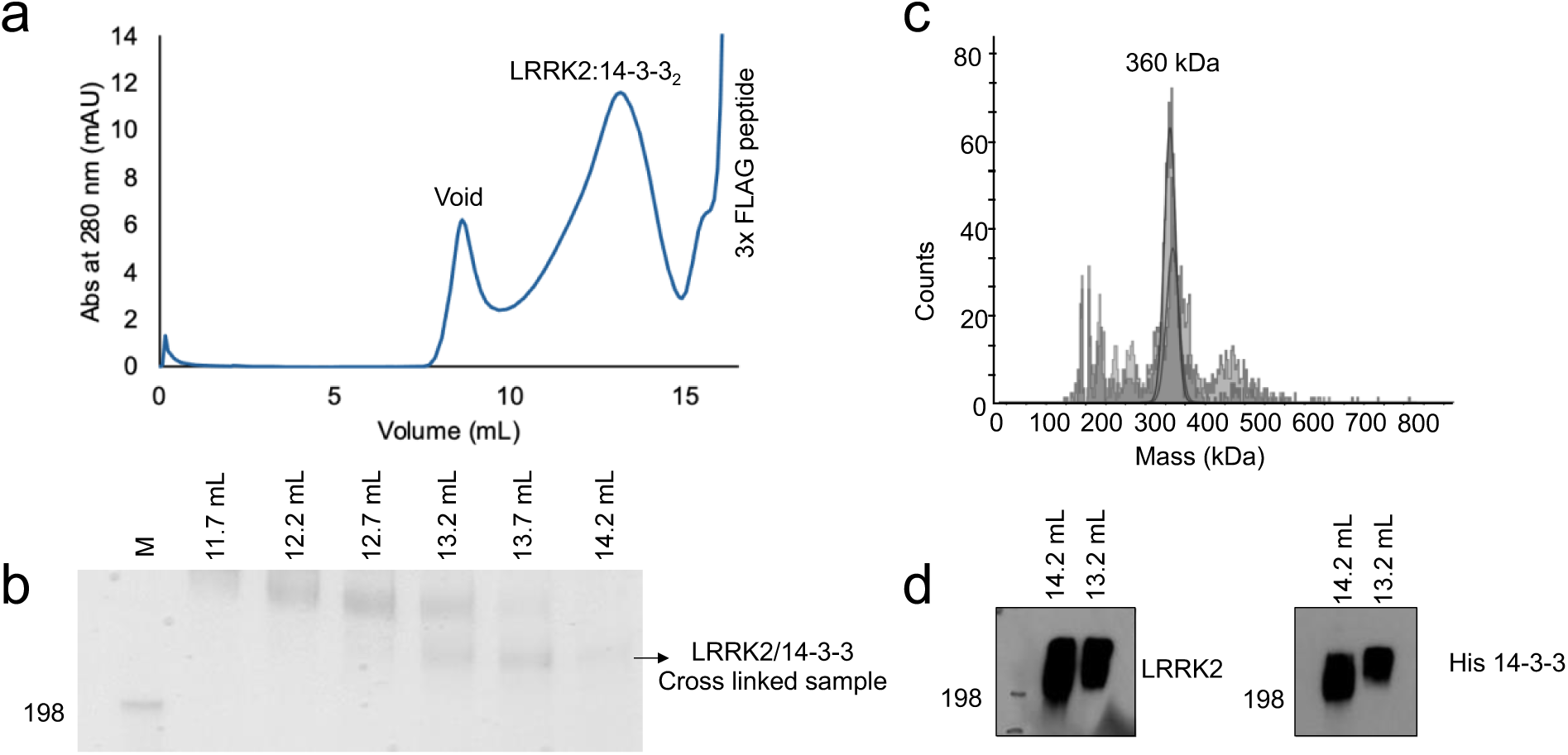
Purification and biochemical characterization of the BS3- cross-linked LRRK2:14-3-3_2_ complex. **a.** Gel filtration chromatogram on a Superose 6 Increase 10/30 column showing the elution profile of the complex. **b.** 4-12% SDS-PAGE gel analysis indicating the presence of the cross-linked cross LRRK2/14-3-3 band. **c.** Mass photometry analysis depicting the mass distribution of the corresponding gel filtration LRRK2:14-3-3_2_ fraction. **d.** Western blot confirming the cross-linking of FLAG- LRRK2 and His-14-3-3 γ, using anti-LRRK2 primary antibody (Abcam cat. ab133474) anti-14-3-3 γ (Abcam cat.137048) were used.

**Supplementary Fig. 5.**
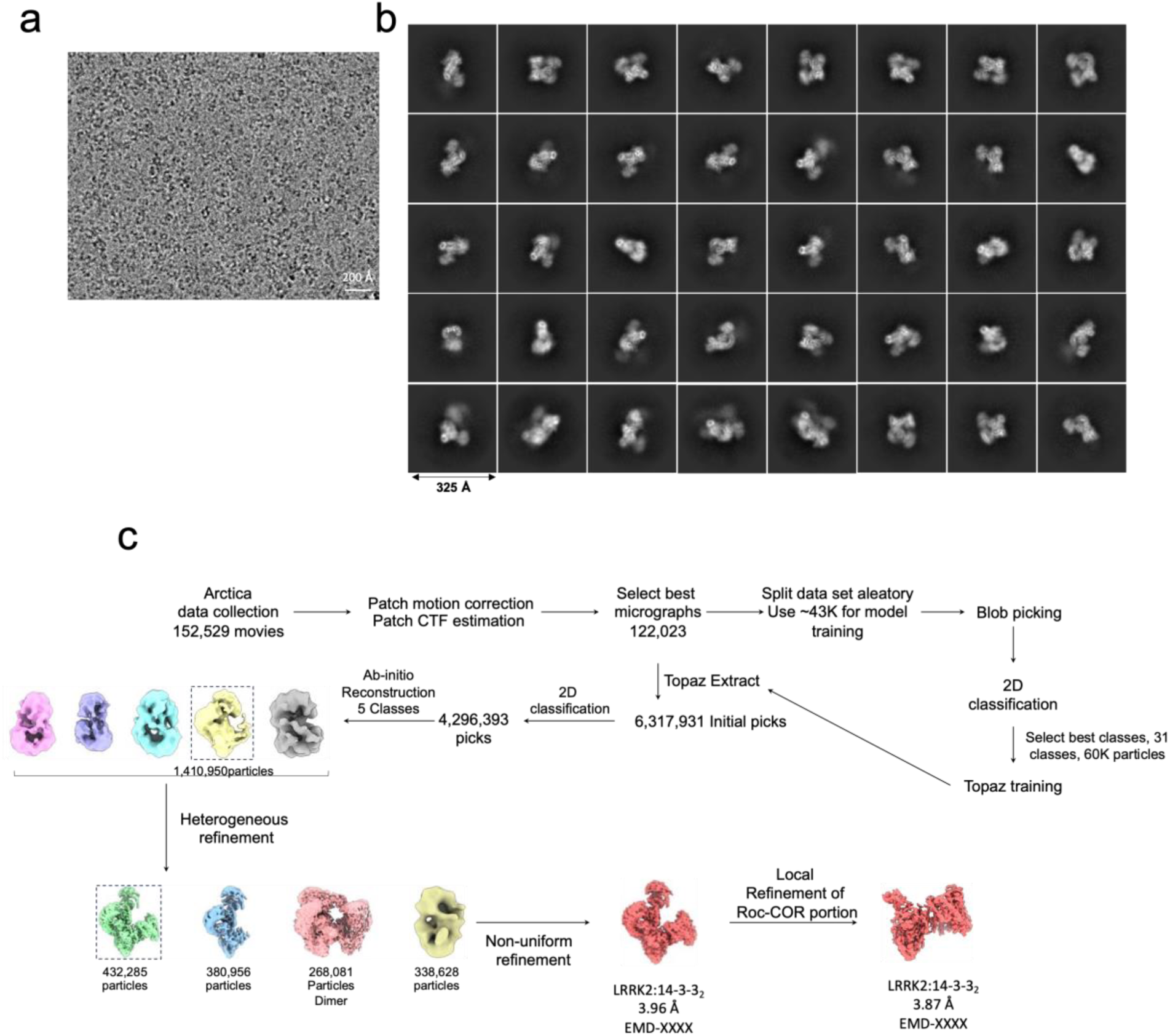
Cryo-EM data collection and processing of the LRRK2:14- 3-3_2_ complex. **a.** Representative cryo-EM image of micrographs **b.** Selected 2D class averages of LRRK2:14-3-3_2_ particles **c.** Schematic of the cryo-EM 3D reconstruction workflow.

**Supplementary Fig. 6.**
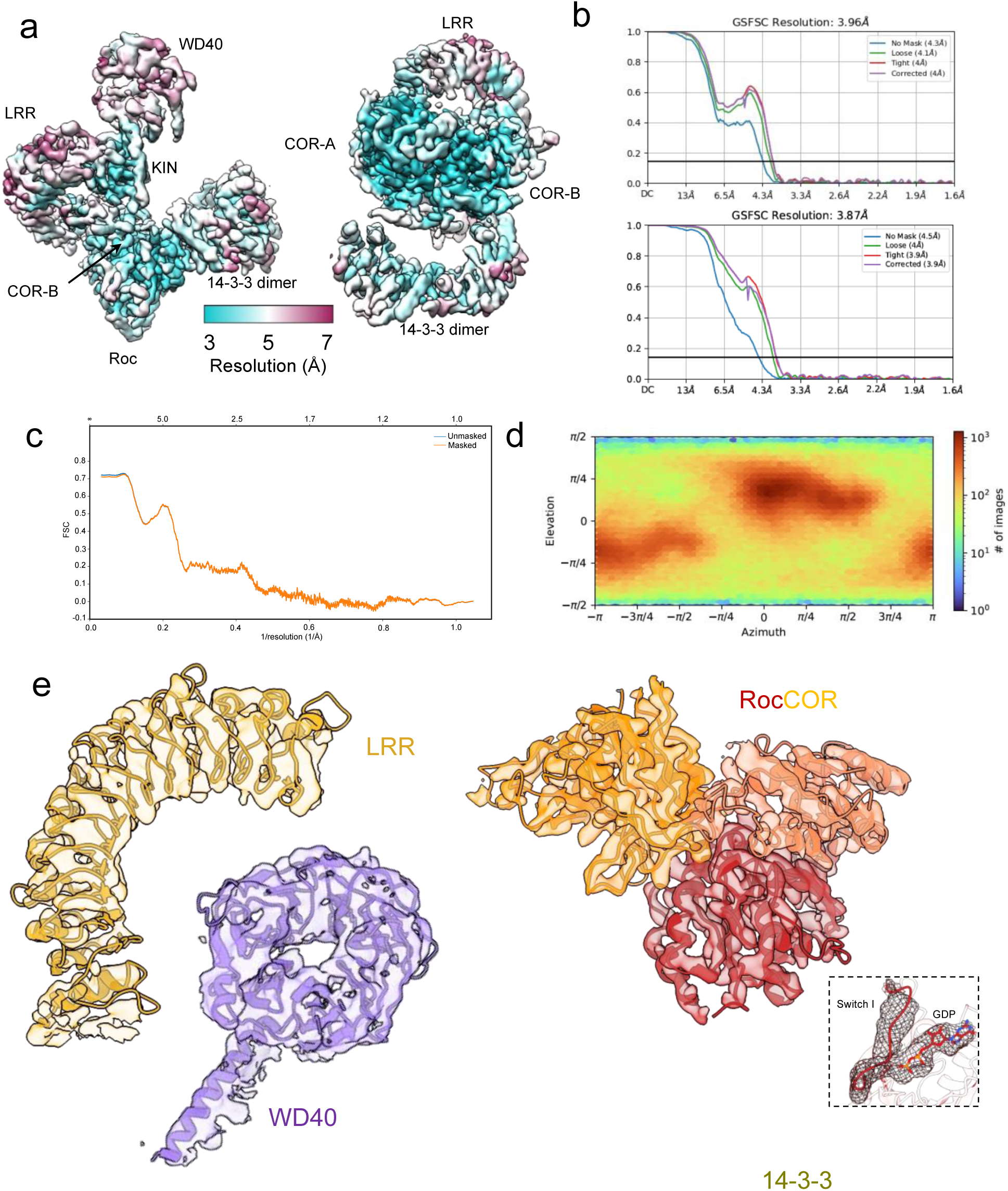
Comprehensive cryo-EM analysis of the LRRK2:14-3-3_2_ complex. **a.** Cryo-EM map displayed with color coding according to local resolution. **b.** Golden-standard Fourier shell correlation (FSC) curve for the resolution assessment of the global (top) and Roc-COR:14-3-3 local (bottom) resolution maps **c.** Map-model FSC curve verifying model fit to the cryo-EM density map **d.** Angular distribution plot of particle orientations for the global non-uniform refinement. **e.** Detailed views of density regions of specific LRRK2 domains and 14-3-3 protein within the complex.

**Supplementary Fig. 7.**
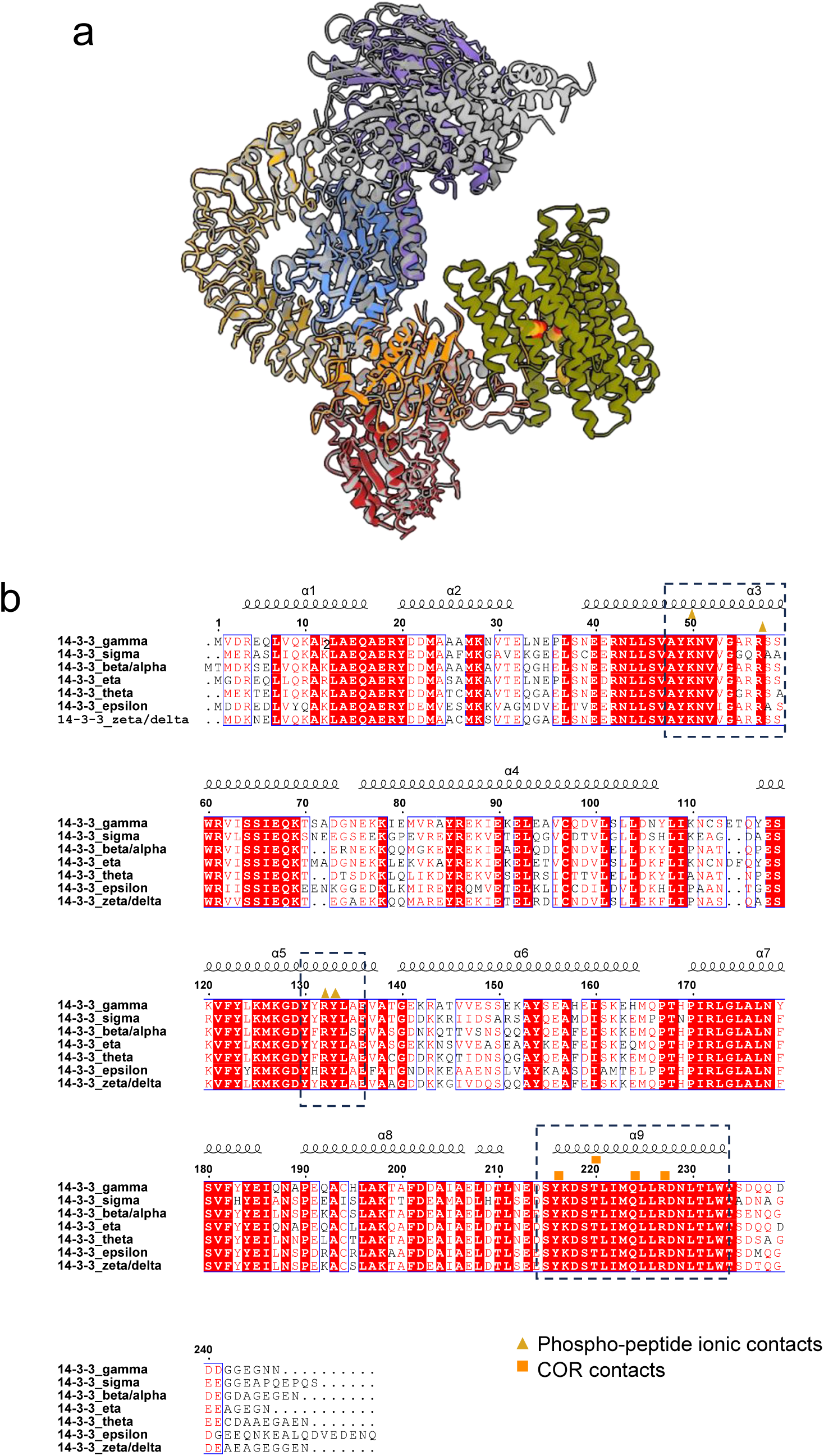
Comparative structural and sequence analysis of LRRK2 interactions with 14-3-3. **a.** Superposition of the LRRK2:14-3-3_2_ complex structure (colored as in Fig. 1) with the previously reported cryo-EM structure of the unliganded monomeric LRRK2 (PDB: 7LHW) shown in grey. **b.** Sequence alignment of human 14- 3-3 isoforms. The secondary structure is indicated above the alignment. Identically conserved residues are shaded in red. Symbols above the alignment indicate 14-3-3 residues involved in the phospho-peptide ionic contacts (golden triangles) and COR contacts (orange squares). Dashed squares highlight these regions.

**Supplementary Fig. 8.**
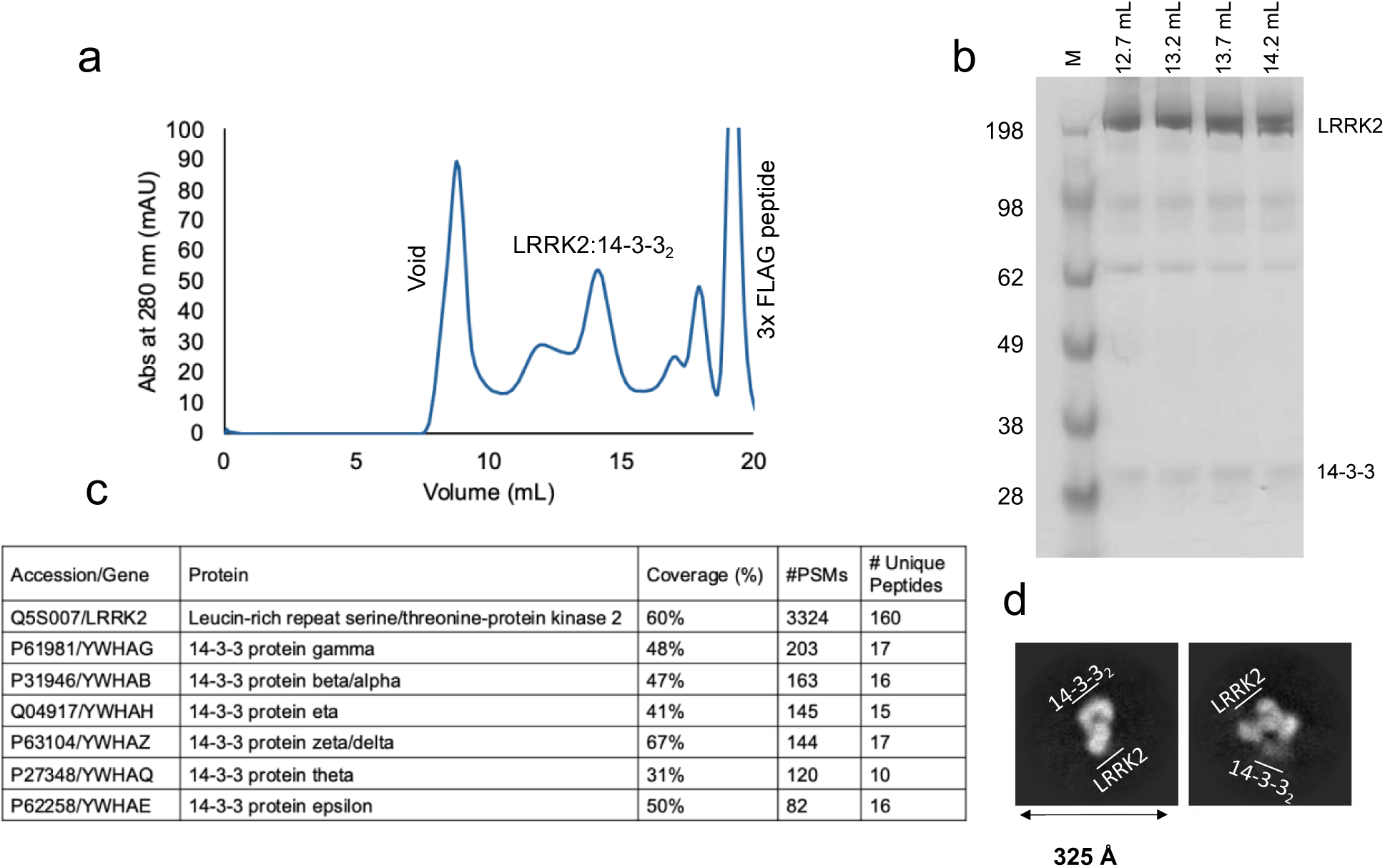
Isolation and characterization of human LRRK2/14-3-3 complex formed in a cellular context. **a.** Gel filtration chromatography of the LRRK2/14-3-3 complex using a Superose 6 Increase 10/30 column. **b.** SDS-PAGE analysis confirms the formation of the complex. **c.** Mass spectrometry validated the complex formation. Number of total (Peptide Spectrum Matches (PSMs) and unique peptides are reported. **d.** Diagnostic cryo-EM depicting representative 2D class averages.

**Supplementary Fig. 9.**
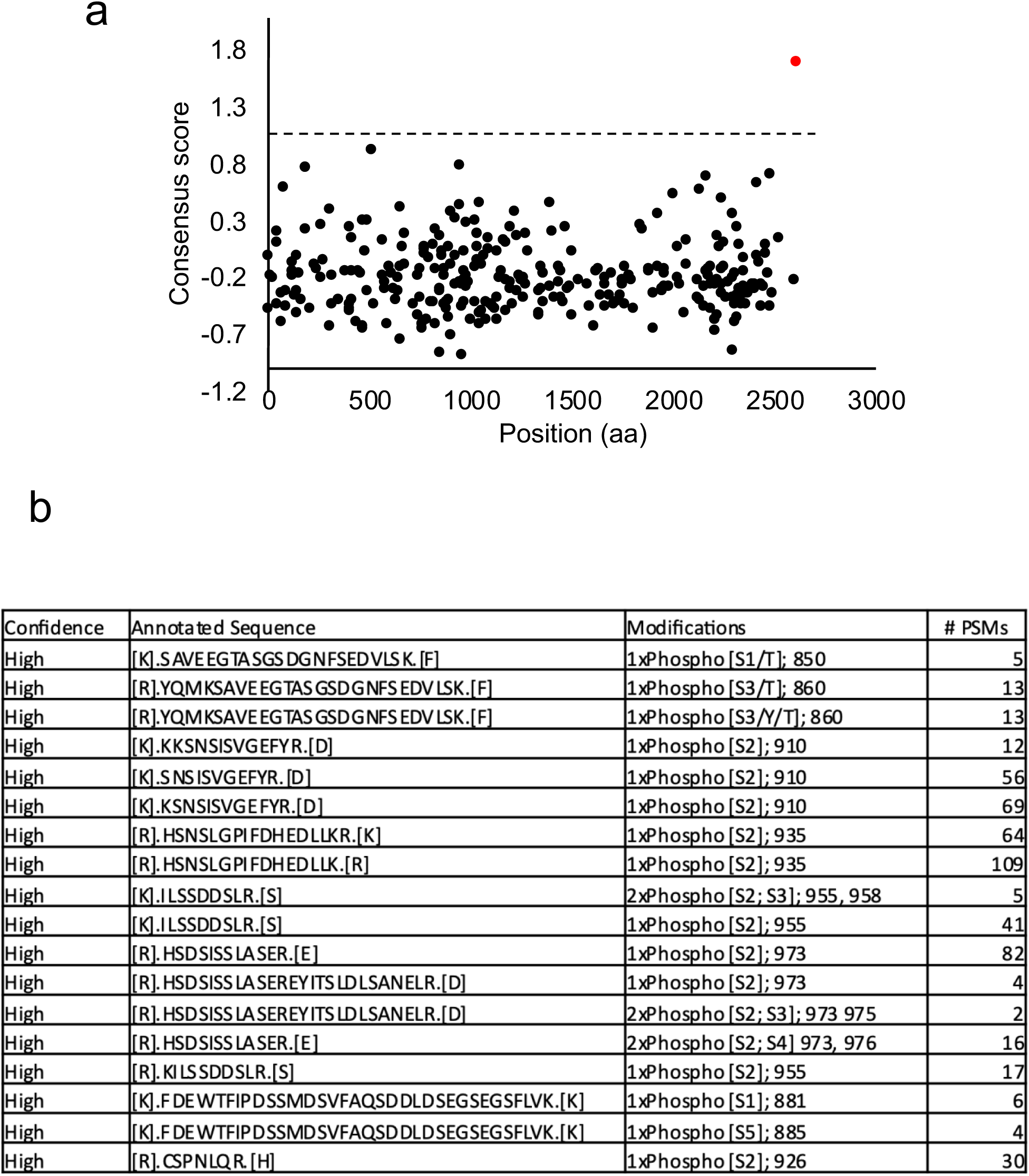
Analysis of putative 14-3-3 binding sites in LRRK2. **a.** Prediction of 14-3-3 consensus binding sites in LRRK2, based on three different classifiers (ANN, PSSM, and SVM) using the 14-3-3-Pred webserver^81^. Data points represent the putative binding sites in the LRRK2 sequence. The arbitrary red point indicates the consensus score for a conventional binding motif (RSXSXP). No outstanding candidates are identified in LRRK2 by the consensus score. **b.** Identification of phospho-sites in purified LRRK2 protein by mass spectrometry, the number of peptides (Peptide Spectrum Matches (PSMs), and confidence in phosphorylation are reported.

**Supplementary Fig. 10.**
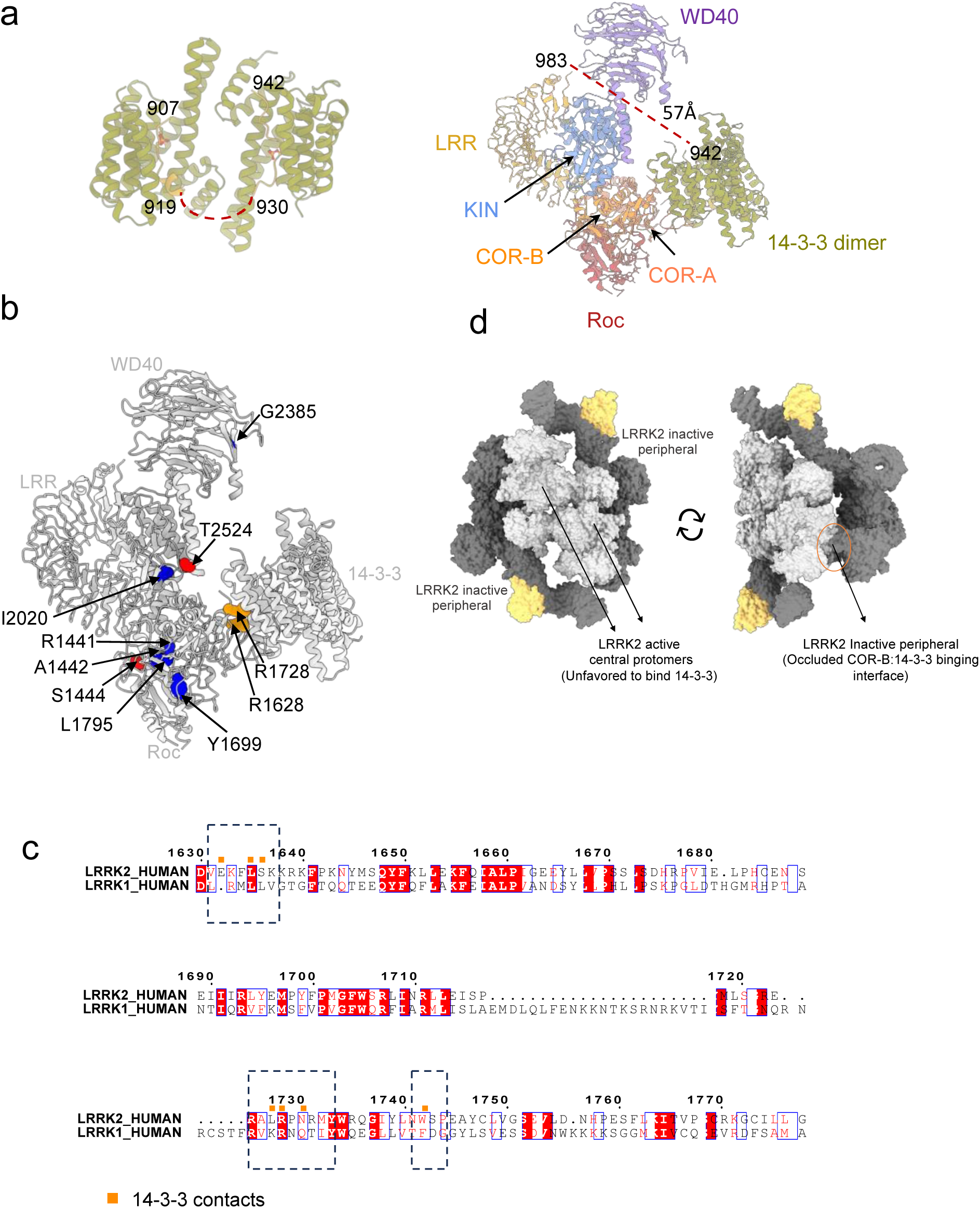
Additional structural analysis and hypothetical binding regions in the LRRK2:14-3-3_2_ complex. **a.** Illustration of missing loop regions connecting the 14-3-3-binding-phoshopeptides with the LRR domain and between these regions are depicted with dashed red lines. **b.** Mapping of additionally proposed 14-3-3 phospho-binding sites from previous studies^77^ in red. PD mutations at (orange) and away from (blue) the 14-3-3 binding interface are also shown. **c.** Unfavourability of the LRRK2/14-3-3 interaction in the LRRK2 tetramer (PDB: 8FO9). The active central LRRK2 protomers are shown in light gray, the inactive peripheral LRRK2 protomers in dark gray, and Rab29 in yellow. **d.** Sequence alignment of the COR domain in human LRRK2 and LRRK1, highlighting identically conserved residues in red. Symbols above the alignment indicate the 14-3-3 contacts (orange squares), with dashed squares highlighting these regions.

**Supplementary Fig. 11.**
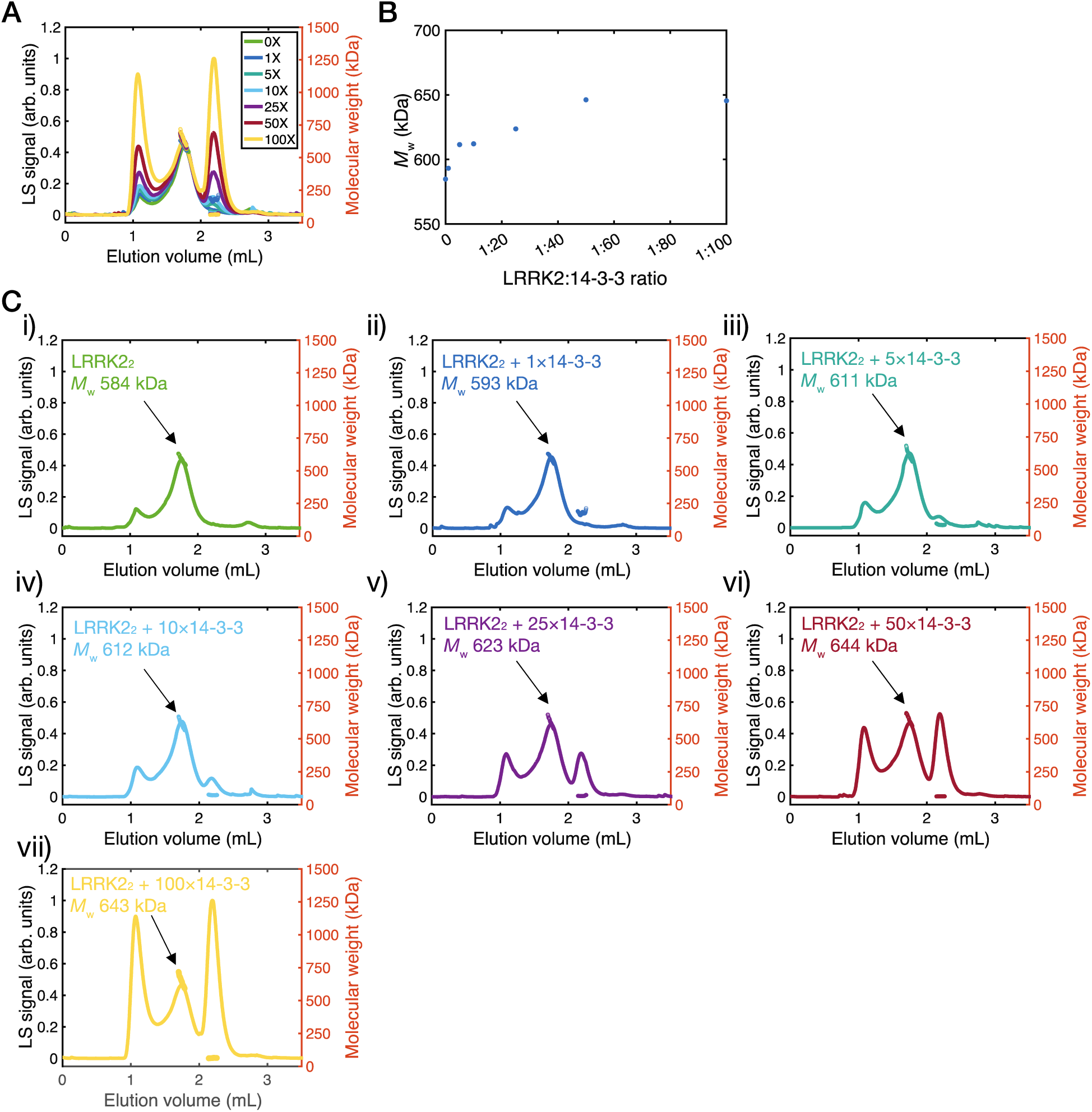
Interaction of LRRK2 dimer with 14-3-3. **a.** Overlay of the MALS chromatograms for the LRRK2 illustrating the effect of increasing concentrations of 14-3-3. **b.** Demonstrating a 14-3-3-concentration dependent increase in LRRK2 mass, consistent with the binding of a single 14-3-3 dimer to a LRRK2 dimer which has a molecular weight of 616 KDa. **c.** Individual chromatograms from panel (a).

**Supplementary Fig. 12.**
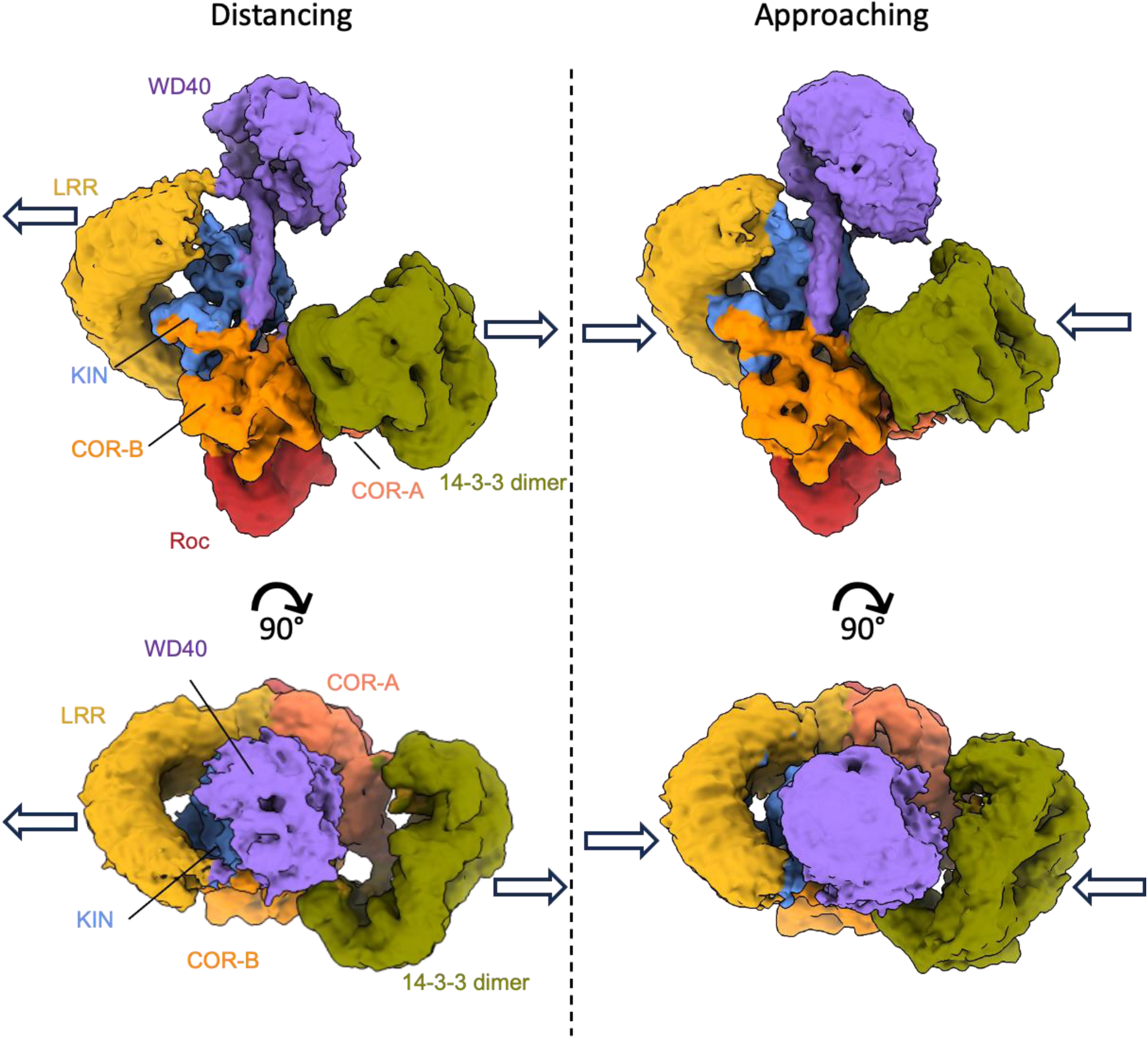
Dynamic Interactions in the LRRK2:14-3-3_2_ complex by 3D variability analysis (3DVA). Analysis demonstrates the synchronized movement of the LRR domain and 14-3-3 dimer relative to the Roc-COR-Kinase-WD40 portion of the protein. Arrows indicate the directions of motion captured by the 3D variability analysis. See also, Supplementary Movie 1. Figures and movie were generated using a filter resolution of 7 Å to resolve and visualize the 3DVA component.

**Supplementary Fig. 13.**
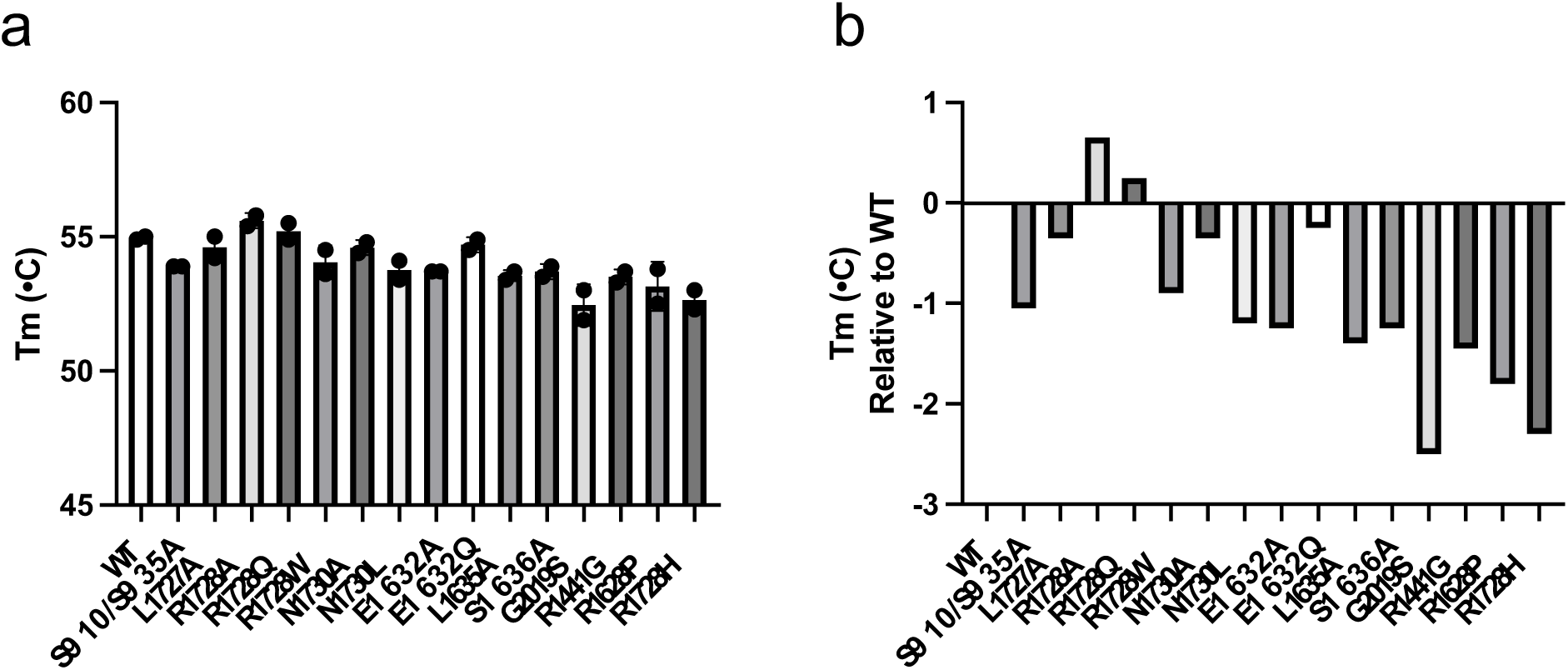
Thermal stability analysis of LRRK2 variants. **a.** Melting temperature (Tm) of LRRK2 variants involving mutations at the LRRK2:14-3-3 interaction interface and other PD related mutations. Error bars represent mean ± SEM from two independent experiments. **b.** Comparison of melting temperatures across all tested LRRK2 variants, plotted relative to WT LRRK2. See Source data file for raw data.

**Supplementary Fig. 14.**
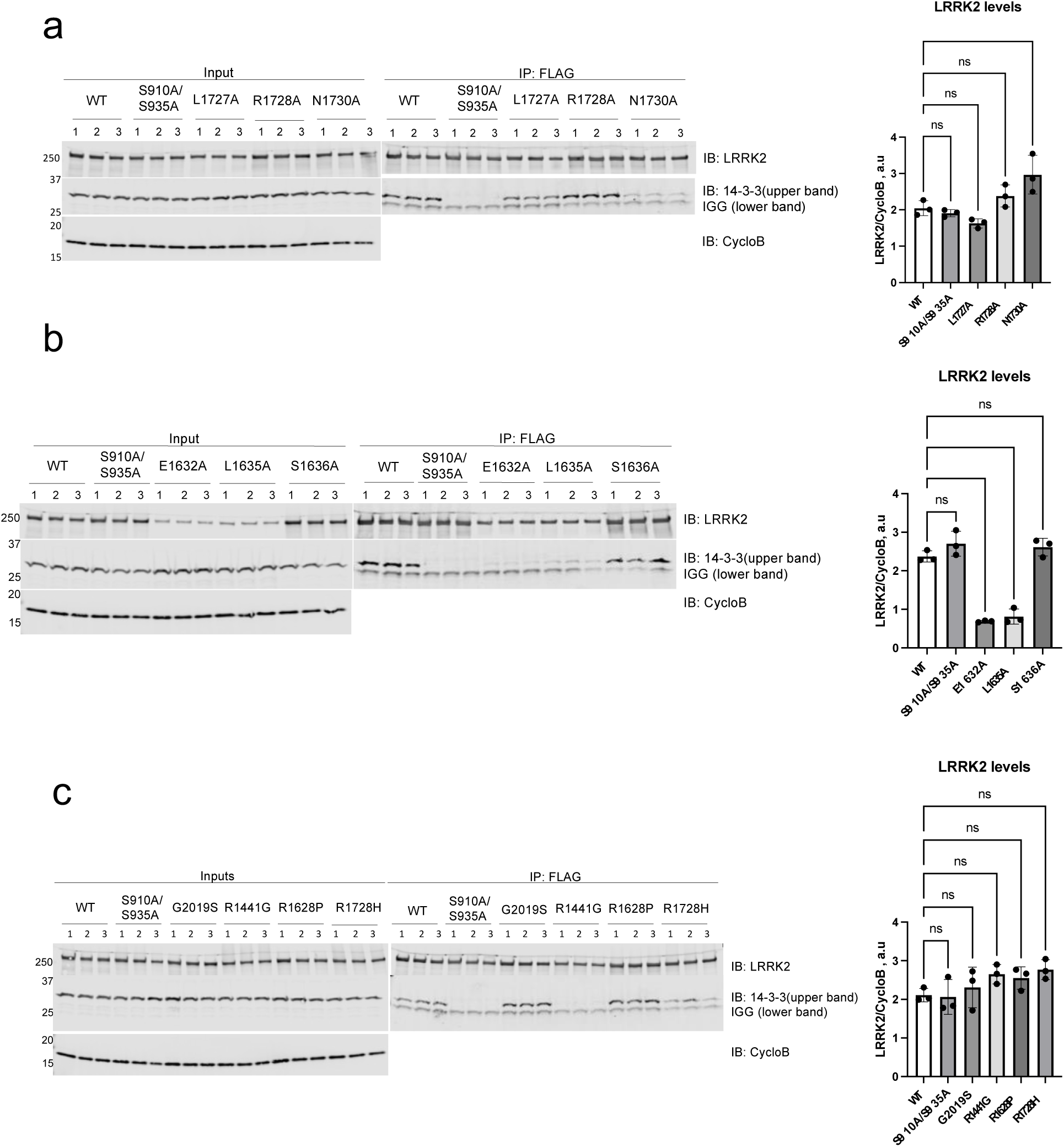
**Representative cropped membrane images for the LRRK2/14-3-3 Co-IP experiments. a-b**: Related to Fig. 2d, c: Related to Fig. 5a,c showing representative data for the results presented in these figure panels. For full uncropped blot images refer to the source data file.

**Supplementary Table 1.**
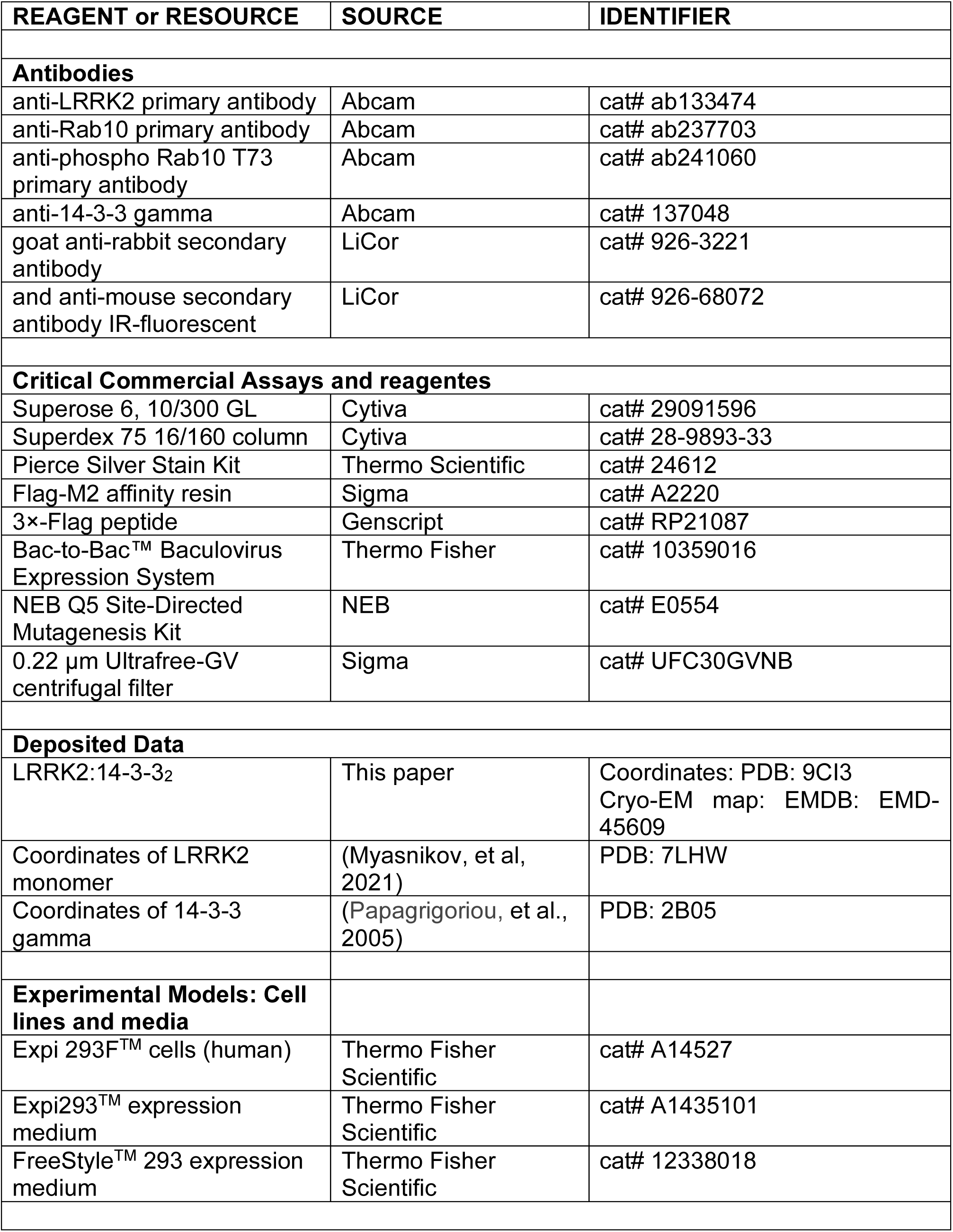
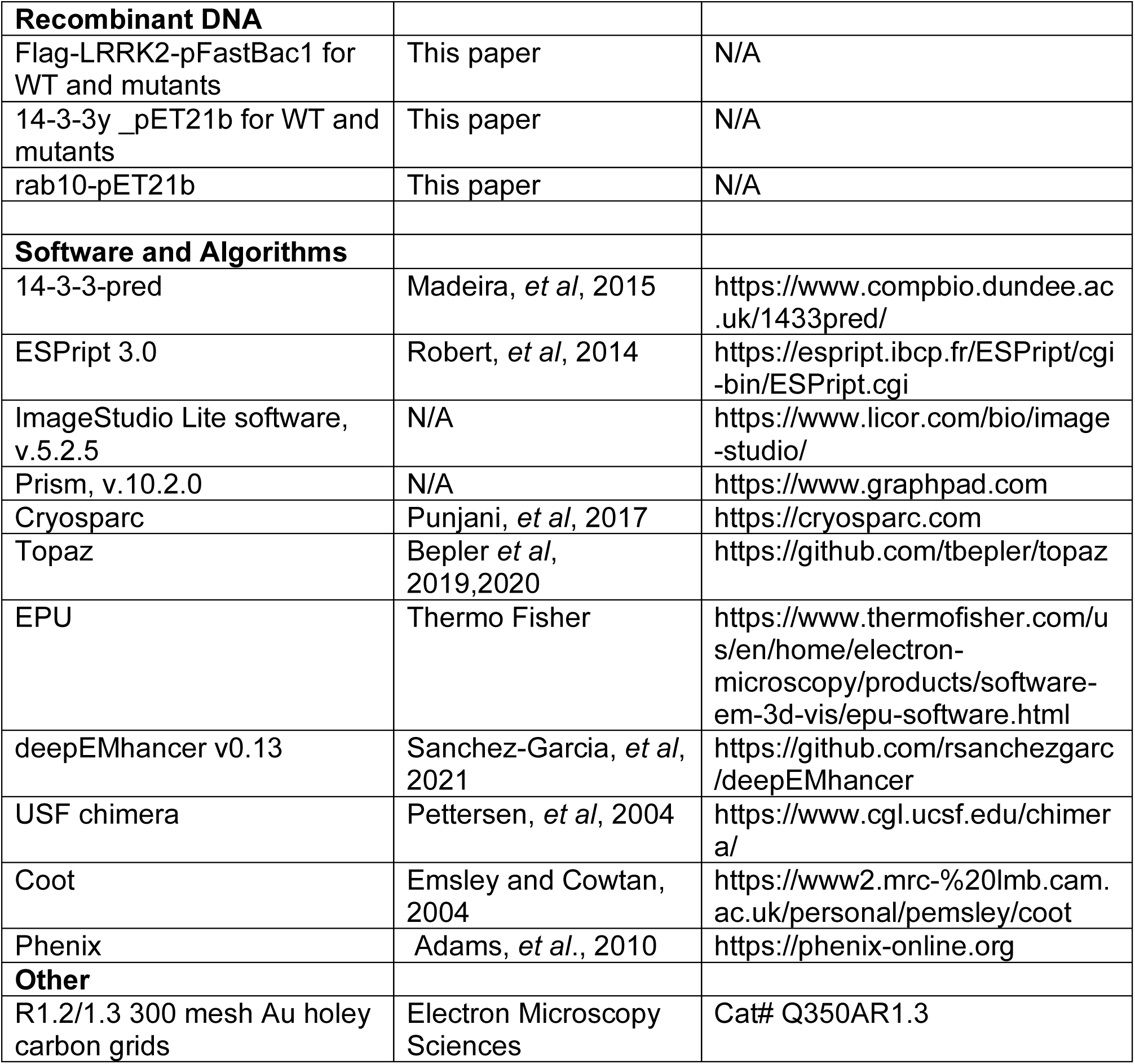
Reagent or Resource Table.

**Supplementary Table 2.** Cryo-EM data collection, Refinement and Validation Statistics.

|  | **LRRK2:14-3-3<sub>2</sub>** |
| --- | --- |
| **Data collection** |  |
| Microscope | Talos Arctica |
| Camera | K3 |
| Magnification | 100k, EFTEM mode |
| Voltage (kV) | 200 |
| Electron exposure (e<sup>-</sup>/Å<sup>2</sup>) | 50 |
| Number of frames collected per micrograph | 50 |
| Energy filter slit width | 20eV |
| Defocus range (μm) | -0.8 to -2.5 |
| Pixel size (Å) | 0.81 |
| Movies (no.) | 152,529 |
| Initial particle images (no.) | 6,317,931 |
| Final particle images (no.) | 432,285 |
| Map resolution (Å) | 3.96 |
| FSC threshold | 0.132 |
| Map sharpening B factor (Å<sup>2</sup>) | 171.6 |
| EMDB code |  |
| **Model building and refinement** |  |
| Initial model used (PDB) | 7LHW, 2B05 |
| Model composition |  |
| Non-hydrogen atoms | 15949 |
| Protein residues | 2002 |
| Protein molecules | 3 |
| Real-space correlation |  |
| CCvolume | 0.58 |
| CCmask | 0.58 |
| Mean B factor (Å<sup>2</sup>) | 55.65 |
| RMS deviations |  |
| Bond lengths (Å) (outliers >4σ) | 0.003 (0) |
| Bond angles (°) (outliers >4σ) | 0.777 (16) |
| Validation |  |
| MOLProbity score | 2.34 |
| Clashscore | 19.16 |
| Rotamer outliers (%) | 1.30 |
| CaBLAM outliers (%) | 0.20 |
| Cβ outliers (%) | 0.00 |
| Ramachandran plot |  |
| Favored (%) | 92.32 |
| Allowed (%) | 7.63 |
| Outliers (%) | 0.05 |
| PDB code | 9CI3 |

## Notes

### Competing Interest Statement

The authors have declared no competing interest.

